# Chemophoresis Engine: Universal Principle of ATPase-driven Cargo Transport

**DOI:** 10.1101/2021.10.15.463834

**Authors:** Takeshi Sugawara, Kunihiko Kaneko

**Affiliations:** Universal Biology Institute, The University of Tokyo, 7-3-1 Hongo, Bunkyo-ku, Tokyo 113-0033, Japan; Center for Complex Systems Biology, Universal Biology Institute, University of Tokyo, 3-8-1 Komaba, Meguro-ku, Tokyo 153-8902, Japan

## Abstract

Cell polarity regulates the orientation of the cytoskeleton members that directs intracellular transport for cargo-like organelles, using chemical gradients sustained by ATP or GTP hydrolysis. However, how cargo transports are directly mediated by chemical gradients remains unknown. We previously proposed a physical mechanism that enables directed movement of cargos, referred to as chemophoresis. According to the mechanism, a cargo with reaction sites is subjected to a chemophoresis force in the direction of the increased concentration. Based on this, we introduce an extended model, the *chemophoresis engine*, as a general mechanism of cargo motion, which transforms chemical free energy into directed motion through the catalytic ATP hydrolysis. We applied the engine to plasmid motion in a parABS system to demonstrate the the self-organization system for directed plasmid movement and pattern dynamics of ParA-ATP concentration, thereby explaining plasmid equi-positioning and pole-to-pole oscillation observed in bacterial cells and *in vitro* experiments. We mathematically show the existence and stability of the plasmid-surfing pattern, which allows the cargo-directed motion through the symmetry-breaking transition of the ParA-ATP spatiotemporal pattern. Finally, based on its generality, we discuss the chemophoresis engine as a universal principle of hydrolysis-driven intracellular transport.

**SIGNIFICANCE:** The formation of organelle/macromolecule patterns depending on chemical concentration under non-equilibrium conditions, first observed during macroscopic morphogenesis (1), has recently been observed at the intracellular level as well, and its relevance as intracellular morphogen has been demonstrated in the case of bacterial cell division. These studies have discussed how cargos maintain positional information provided by chemical concentration gradients/localization. However, how cargo transports are directly mediated by chemical gradients remains unknown. Based on the previously proposed mechanism of chemotaxis-like behavior of cargos (referred to as chemophoresis, (2, 3)), we introduce a *chemophoresis engine* as a physicochemical mechanism of cargo motion, which transforms chemical free energy to directed motion. The engine is based on the chemophoresis force to make cargoes move in the direction of the increasing ATPase(-ATP) concentration and an enhanced catalytic ATPase hydrolysis at the positions of the cargoes. Applying the engine to ATPase-driven movement of plasmid-DNAs in bacterial cells, we constructed a mathematical model to demonstrate the self-organization for directed plasmid motion and pattern dynamics of ATPase concentration, as is consistent with *in vitro* and *in vivo* experiments. We propose that this chemophoresis engine works as a universal principle of hydrolysis-driven intracellular transport.

## INTRODUCTION

Cell polarity regulates the direction of intracellular transport for cargos, such as organelles and macromolecules, by taking advantage of chemical gradients sustained with the aid of ATP or GTP hydrolysis. For example, it is well known that eukaryotic cell polarity factors, such as Rho, GTPase, and Cdc42, regulate the orientation of cytoskeleton members so that molecular motors can carry cargo directionally on the cytoskeleton, contributing to cell movement (4), cell growth (5), and axon guidance (6). Although transport by the cytoskeleton is important, transport directly mediated by chemical gradients, if possible, should be of importance as a general mechanism for cargo transport as well, which we refer to as cargo chemotaxis.

A bacterial parABS system (7–18) is a good candidate for cargo chemotaxis. It is the most ubiquitous bacterial polarity factor that regulates the separation of bacterial chromosome/plasmids into daughter cells by organizing their regular positioning along the cell axis (7–18). Generally, it consists of three components as follows: The DNA binding protein ParB, ATPase, ParA, and the centromere-like site *parS*. ParB binds *parS*, spreads along the DNA, and forms a large partition complex (PC) around *parS*. ATP-bound ParA (ParA-ATP) can nonspecifically bind to DNA and interact with ParB-*parS* PC. Abundant ParA-ATP molecules are distributed on a nucleoid in a host cell. Their mobility is strongly restricted so that they are not homogenously distributed in the cell, thus enabling a sustained concentration gradient even within a micron-sized cell (14–16, 19–25). Recent reports have suggested that ParA-ATP gradient/localization can drive a *parS* site formation on a host genome/plasmid in the direction of the increased concentration, which can be a major candidate mechanism for plasmid partitioning and chromosome segregation (2, 3, 24–40).

ParA ATPase is an evolutionarily conserved protein which has many homologs (41–44). Representative examples of its family are McdA/McdB ATPases controlling equidistribution of carboxysomes along a long cell axis in cyanobacteria (45–47), ParC/PpfA ATPases that regulate intracellular positions of chemotaxis protein clusters (48–50), MipZ ATPase that coordinates chromosome segregation in cell division (43, 51, 52), and MinD ATPase that determines a cell division plane (53–55). These ATPase homologs, as well as ParA, work through a common mechanism essential to their function: Hydrolysis of ATPase X by a partner protein B; X − ATP + B ⇄ Y → X + ADP + B. By taking advantage of the free energy released by the reaction, a spatiotemporal pattern of the corresponding ATPase emerges (23, 43, 53–59), and cargo positions are coordinated (45–50). One of the most renowned intracellular patterning systems is the MinCDE system that self-organizes the pole-to-pole oscillation of MinD, leading to the formation of a cell division plane at the cell center, upon stimulation of MinD ATPase activity induced by MinE at the inner cell membrane (54, 55). In contrast, in the *in vitro* reconstitution of the Min system, traveling waves of MinD were observed (55–59).

Similar to the Min system, the pole-to-pole oscillation of ParA (15, 24, 60–64) also emerged in the parABS system through the stimulation of ParA ATPase activity by ParB on the PC (14–16, 23–25). Interestingly, a plasmid chases a ParA focus, following its oscillatory movement along the long host-cell axis (24), leading to oscillatory motion. In a recent *in vitro* experiment mimicking a parABS system, Vecchiarelli et al. elegantly demonstrated the formation of directed motion of a cargo corresponding to a plasmid, referred to as “cargo surfing on ParA-ATP traveling wave” (27–29). Hence, for both Min and Par systems, the emergence of traveling waves and the pole-to-pole oscillation of ATPase have been reported. The mechanism driving the plasmid motion, however, remains elusive (19), whereas the pattern dynamics of the Min system can be described by well-defined reaction-diffusion equations (55–57, 65–69).

Previously, we proposed a mechano-chemical coupling mechanism that enables directed movement of cargos, referred to as chemophoresis (2, 3). According to this mechanism, a macroscopic object with reaction sites on its surface is subjected to a thermodynamic force along an increasing concentration gradient. Cargo transport is possible via the chemophoresis force, (2, 3), and the possible role of the chemophoresis force in the separation dynamics of bacterial plasmids was discussed previously. By combining the plasmid motion driven by the chemophoresis force with a reaction-diffusion (RD) equation, we demonstrated that regular positioning of plasmids is possible in a parABS system under ParA-ATP hydrolysis stimulated by ParB (17, 25, 61–64). To date, however, spontaneous directed motion or pole-to-pole oscillation of plasmids (24, 61–64) has not been discussed in Ref (3), as was demonstrated by Vecchiarelli et al. (27–29) and theoretical studies (33, 35–38).

In the present study, we extend our previous model and propose a *chemophoresis engine* as a general mechanism of cargo motion, which transforms chemical energy into directed motion via self-organization of the traveling wave, and then apply it to the plasmid motion in a parABS system. In the previous study, the plasmid was assumed to be a point particle, where static equi-positioning and symmetric ParA-ATP distribution were robustly maintained (3). However, such model with a zero-size limit is unrealistic, considering intracellular dynamics or reconstructing *in vitro* experiments performed by (27). Here, by considering the finite size of plasmids explicitly, we show that organization of directed motion is possible via spontaneous symmetry breaking in the ParA-ATP pattern. We then recapitulate plasmid positioning to better describe the spatiotemporal profiles of ParA-ATP concentration and movement of plasmids. Actually, in the model presented here, the net chemophoresis force acts on the plasmid (PC) through the concentration difference between its ends, which is self-sustained by the high ATP hydrolysis stimulation. This self-driven mechanism leads to the directed motion of plasmids, as well as their equi-positioning (17, 25, 61, 64), and pole-to-pole oscillation as observed in bacterial cells and *in vitro* experiments (24, 61–64). We mathematically show the existence and stability of the plasmid-surfing pattern, which allows cargo-directed motion through the symmetry-breaking transition of the ParA-ATP spatiotemporal pattern.

## METHODS

### Numerical methods for solving evolutionary equation, self-consistent equation, and eigenvalue equation

Evolutionary equations, Eq. 3 and 4 were computationally solved as a hybrid simulation between reaction-diffusion equation and Langevin equation. Euler scheme for Eq. 3 and Euler-Maruyama scheme for Eq. 4 were used as numerical algorithms. The Hill coefficient *m* were fixed through the simulation as *m* = 1. Following parameters were used in the simulation except for Fig 2D: *l*_*b*_ = 0.2, *N* = 40. Real-valued self-consistent equation, Eq S9 and complex-valued eigenvalue equation S19 (Supporting Materials and Methods) were solved by using Newton-Raphson method.

## RESULTS

### Chemophoresis force

First, we briefly reviewed the chemophoresis force, a thermodynamic force acting on the cargo in the direction of the increased concentration of a chemical that can be bound on the cargo (See Appendix A and B for details.) We considered that a cargo was placed and moving in a *d*-dimensional space ***r*** *∈****R***^*d*^. The cargo had *N* molecular sites B, on each of which *m* molecules of chemical X was bound to form a complex Y at position ***r*** = ***ξ***. At each site, the reaction mX **(*ξ*)** +B (⇄) + Y occurred and was at chemical equilibrium. If a spatial gradient of chemical concentration X exists, the cargo is thermodynamically driven in the direction of the decreased free energy or the increased chemical potential of X (2, 3). Here, such a gradient of the chemical potential *µ*(***r***) was assumed to be sustained externally through several active processes, supported by spatially distributed chemical gradients. We referred to the phenomenon as *chemophoresis*. The formula of the chemophoresis force was:

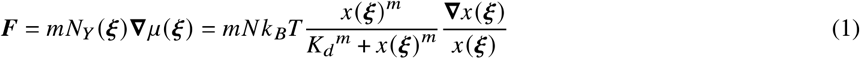

Here, 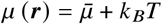 ln *x* (***r***), where *x* (***r***) is the concentration of X,*K*_*d*_ is the dissociation constant, and *m* is the number of binding molecules corresponding to the Hill coefficient of the reaction. With this chemophoresis force, the cargo moved in the direction such that the concentration *x* (***r***) increased even under thermal fluctuation ((2, 3), Appendix B). For the force to work, the reaction mX B + ⇄Y was required to reach chemical equilibrium fast enough for cargo motion. Therefore, we showed that the chemophoresis force was one of the fundamental thermodynamic forces driven by physicochemical fields. Note that the force had an entropic origin from the viewpoint of statistical mechanics. See also Ref. (3) for details of the derivation from the viewpoint of thermodynamics and statistical mechanics.

To understand the origin of chemophoresis, it should be noted that microscopic binding events of X do not directly generate the force. Rather, the force works in the direction of larger frequency of the binding events (or larger time fraction of binding states) that was realized in the spatial location with a larger concentration of molecules in a chemical bath. The chemical gradient biases the binding frequency of X in a space-dependent manner. In other words, chemophoresis is driven by general thermodynamic force as a result of the free-energy (entropy) difference. It can also be derived by coarse-graining microscopic processes (Appendix A and B), whereas the macroscopic derivation implies its generality independent of specific microscopic models (35, 36). On the other hand, both the macroscopic (thermodynamic) and microscopic (statistical physics) theories are equivalent to each other, in that the force is generated with the aid of spatial asymmetry of molecule numbers bound on the bead, if its radius is finite. Further, for the chemophoresis force to act, X molecules do not necessarily have to bind cooperatively to the bead (as in the case of *m* = 1); if the concentration gradient of bound molecules is generated, the resultant free energy difference between its ends leads to the net chemophoresis force.

#### Chemophoresis engine

We then applied the chemophoresis formula to plasmid motion. As the reaction on the cargo consumed chemical X, its concentration changed; therefore, studied its RD equation. It was introduced for ParA-ATP ((2, 3), Appendix B), which recapitulates the central- and equi-positioning of plasmids (2, 3). It was also adopted successfully to explain the directed movement of beads in an *in vitro* experiment by Vecchiarelli et al. (29). We considered a plasmid *i* (1 ≤ *i* ≤ *M*) placed into and moving in a *d*-dimensional space ***r*** *∈* ***R***^*d*^ (*d*= 1 or 2) (Fig. 1 A). ParA-ATP bound to a PC on plasmid *i* at position ***r*** = ***ξ*** *m* ParA-ATP molecules interacted with ParB, which stimulated ParA ATPase activity at a catalytic rate *k*; *N* ParB molecules were assumed to be recruited to each PC at ***r*** = ***ξ*** Because ParA could not bind PC when it was not combined with ATP, free ParA products were released from the PC immediately after ATP hydrolysis. Thus the reaction was presented as follows:

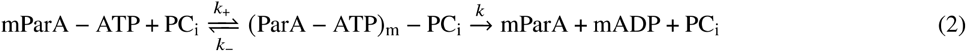

Through this reaction on PC_i_ at ***r*** = ***ξ***, each plasmid acted as a sink for ParA-ATP and induced a concentration gradient of this protein. In the early model, the size of plasmids was assumed to be zero ((2, 3), Appendix B). However, the model is still too unphysical to better reconstruct the movement of plasmids with a finite size in bacterial cells as well as that of micro-sized beads in *in vitro* experiments (27). To better describe spatiotemporal profiles of ParA-ATP concentration and directed movement of the plasmids, we considered plasmids as spheres with a radius of *l*_*b*_ (Appendix B). Denoting the normalized concentration of ParA-ATP as *u*(***r***), the normalized RD equation was written as follows (see Appendix B for its derivation):

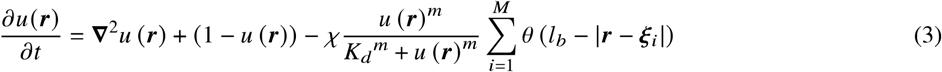

where the first and second terms represent the diffusion of ParA-ATP and its chemical exchange at a normalized constant rate with the cytoplasmic reservoir (denoted by its normalized concentration), respectively. The last term denotes the inhibition by ParB on the *M* PCs. *K*_*d*_ is the normalized dissociation constant of the reaction mX + B ⇄ Y, and *m* is the Hill coefficient. *V* is the *d*-dimensional volume of the bead with a radius of *l*_*b*_. *χ* = *kN*/*V* is a maximum rate for ParA-ATP hydrolysis by ParB on each PC. Furthermore, *θ* (***r***) is a step function representing the space each PC occupies to describe the hydrolysis reaction space. Only within | ***r*** − ***ξ***_***i***_ | < *l*_*b*_, the reaction occurred. Without the last term (if *χ*= 0), *u*(***r***) reached a homogenous equilibrium state, *u*(***r***) = 1. In contrast, the normalized equations of motion for plasmids were represented as follows:

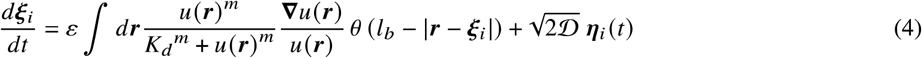

with thermal noise ⟨ ***η*** _***i***_ (*t*) ⟩ = 0 and ⟨ ***η*** _***i***_ (*t*) ***η*** _***j***_ (*t′*) ⟩= 2*dδ*_*ij*_ *δ* (*t* − *t*^′^), and ε = 𝒟 *N/V*. Here, 𝒟= *D*_ξ_ / *D*_*u*_ is the relative diffusion coefficient of the plasmid to that of ParA-ATP (see Appendix B for details). The parameters to be assigned to Eq. 3 and 4 are *K*_*d*_, *l*_*b*_, *m, M, N*, 𝒟, *k*, and the system size *L*(= cell length).

**Figure 1:**
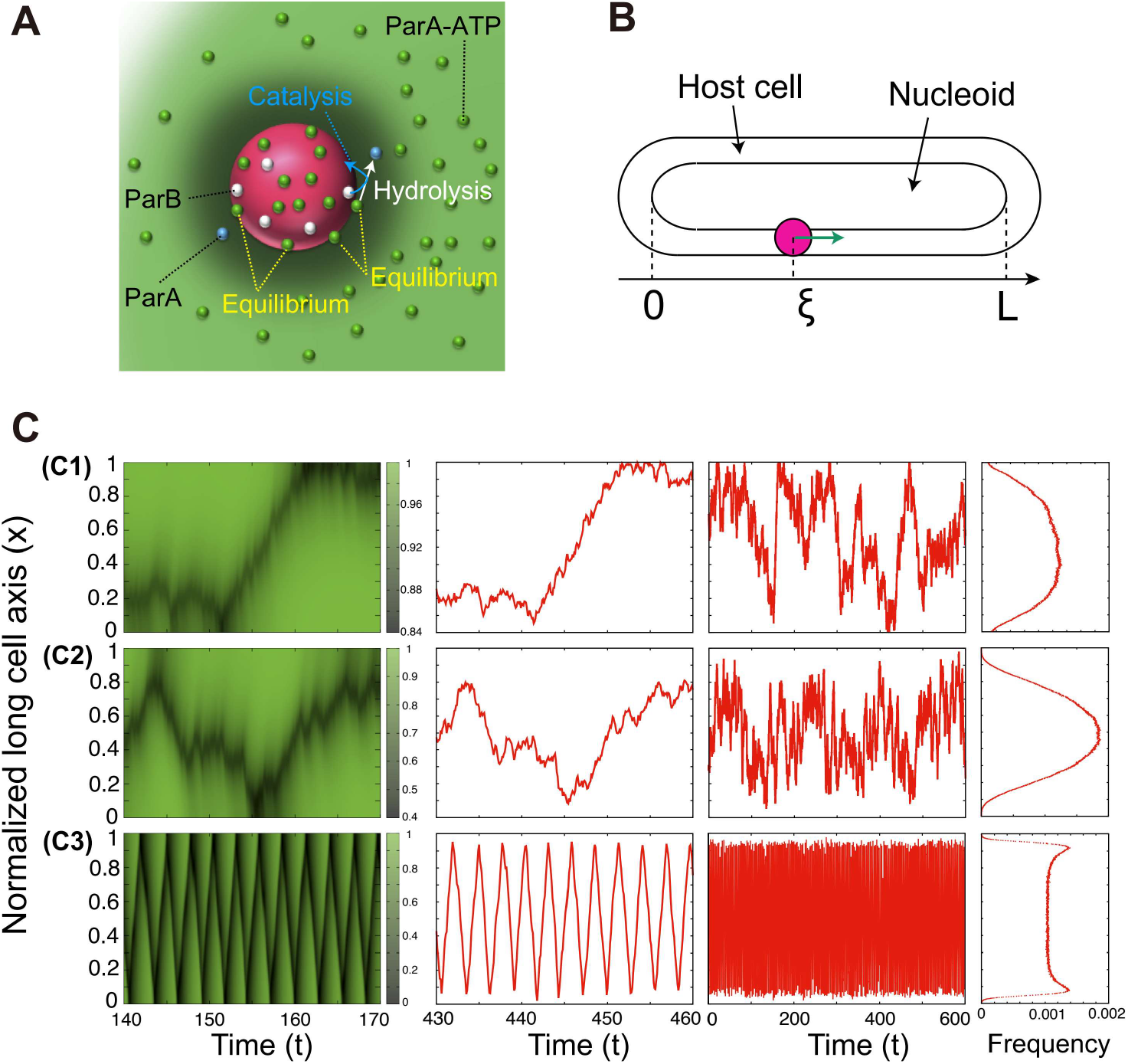
Chemophoresis engine can recapitulate equi-positioning, directed movement, and pole-to-pole oscillation. (A) Schematic representation of the chemophoresis engine. A plasmid moves in a *d*-dimensional space *r ∈* ***R***^*d*^ (*d* = 1 or 2). ParA-ATP (green sphere) binds a partition complex (PC, magenta sphere) on the plasmid at position *r* = *ξ*_*i*_. ParA-ATP molecules interact with ParB molecules (white spheres), which stimulate ParA ATPase activity at a catalytic rate. Because ParA cannot bind PC when it is not combined with ATP, free ParA products (blue sphere) are released from the PC immediately after ATP hydrolysis. Through this reaction on PC_i_ at *r* = *ξ*_*i*_., each plasmid acts as a sink for ParA-ATP and induces a concentration gradient of this protein. (B) One-dimensional case, on a nucleoid matrix along the long cell axis where a plasmid *i* (1 ≤ *i* ≤ *M*) is positioned at *r* = *ξ*_*i*_. ∈ [0,*L*](C) The dynamics change among thermal motion, steady center-positioning, and directed movement followed by oscillatory mode as *χ* increases among *χ* = 0.5 (C1), *χ* = 2.5 (C2), and *χ* = 10 (C3) (two inner figures). (C1) The plasmid slightly tends to be localized at the cell center but it is still dominated by thermal fluctuations for *M* = 1 and *χ* := *kN/V* = 0.5. (C2) It is stably localized at the cell center for *M* = 1 and *χ* = 2.5, and (C3) it shows directed movement, reflection at the end walls, and pole-to-pole oscillation for *M* = 1 and *χ* = 10. The corresponding ParA-ATP pattern dynamics also change among stochastic, steady center-positioning, and oscillatory waves (left). The oscillatory behavior of plasmids does not disrupt time-averaged center-positioning, but steady center-positioning of plasmids are sustained (Compare (C2) right and (C3), right). *K*_*d*_ = 0.1, ε = 5, and *L*= 5. The distributions (right) were generated using 10^7^ samples over 10^5^ time step.

**Figure 2:**
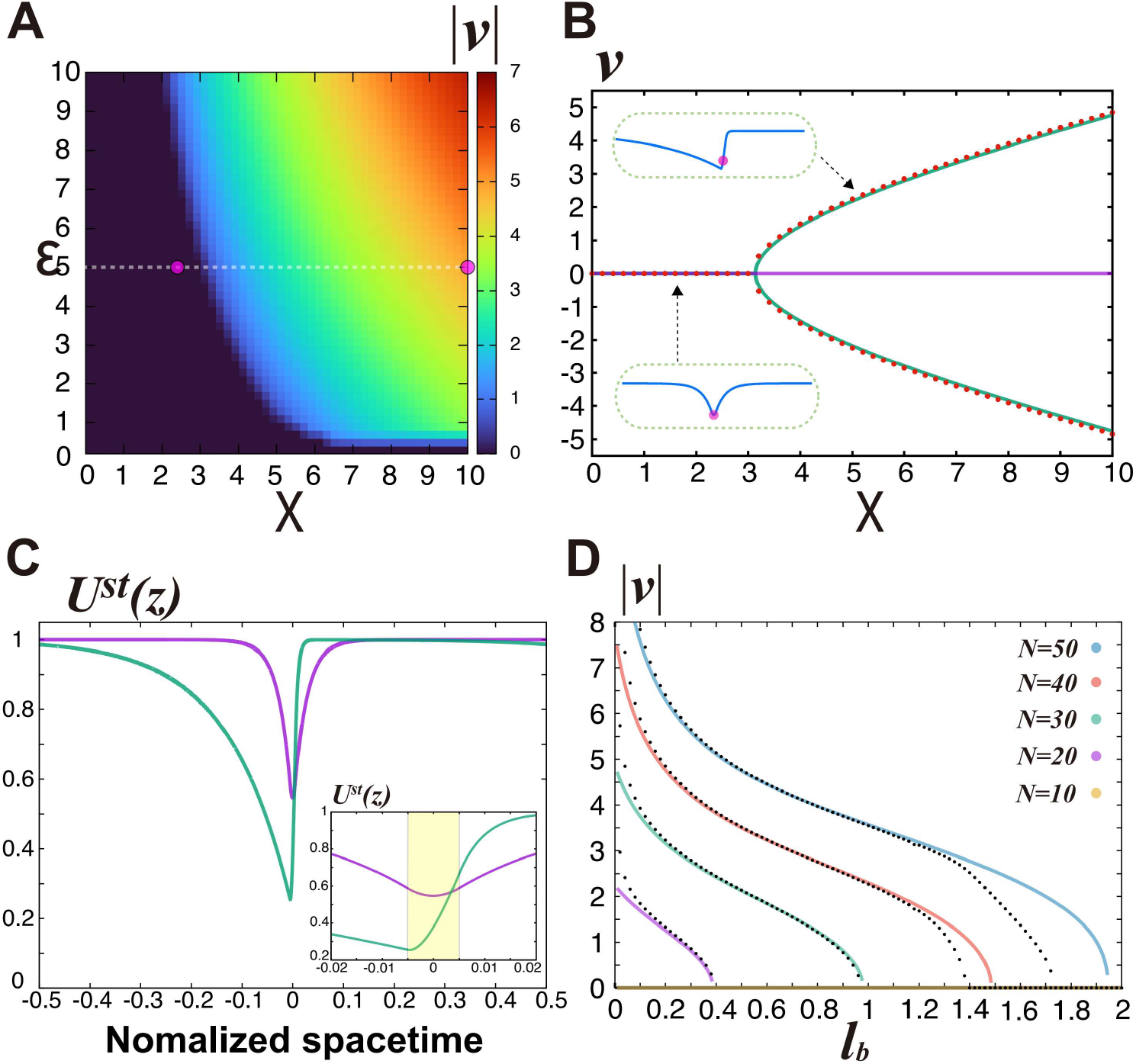
Chemophoresis engine mathematically validates plasmid surfing on the traveling wave of ParA-ATP. (A) Steady velocity (*v*) profile of plasmid movement for 0 < *χ* < 10 and 0 < ϵ < 10 without thermal fluctuations. The plasmid starts moving above a critical curve on the *χ ϵ* plane. The white dotted line shows a parameter region in Fig. 2B. The magenta dots at (*χ, ϵ)* = (2.5, 5), (10, 5)corresponds to parameter values for the steady solutions shown in Fig. 2C.*K*_*d*_ = 0.001, *N* = 40, and *L* = 40. (B) Relationship between *v* and *χ* in analytical (green for *v* ≠ 0 and purple solid line for *v* = 0) for Eq. S3 and S4, and simulated (red dots) solutions for Eq. 3 and 4 (Supporting Materials and Methods). Solutions for directed movement (|*v*| > 0) emerge at *χ* = *χ* _*c*_ ∼ 3.1 as a result of a supercritical pitchfork bifurcation, whereas a solution for localization (*v* = 0) exists over 0 ≤ *χ* ≤ 10.*K*_*d*_ = 0.001, *N* = 40, and *L* = 40. (C) Analytical solutions of the ParA-ATP pattern *U*^*st*^ (*z*) in localization (purple) at *χ* = 2.5 and directed movement (green) at *χ* = 10 for Eq. S6 and S7 (Supporting Materials and Methods). The inset figure shows an enlarged view of *U*^*st*^ (*z*) for *z* /*L∈* [− 0.02 : 0.02]. The plasmid location is fixed at the origin (*z* = 0) on the space-time coordinates. *k*_*d*_ = 0.001, *N* = 40, and *L*= 40. (D) *l*_*b*_ dependency of the directed motion of the plasmid for the analytical solution Eq. (S7 - S9) (solid lines) and numerical result calculated from Eq. 3 and 4 (black dots). For each different value of *N* (= 20, 30, 40, 50), the velocity of the directed motion (*v* ≠ 0) is monotonously decreased with *l*_*b*_, and an inverse pitchfork bifurcation occurs at a critical value of *l*_*b*_, resulting in the only solution with *v* = 0. The numerical results deviate from the analytical ones in the ranges of small and large *l*_*b*_, suggesting the breakdown of the approximation *u*(*x*) ≫*K*_*d*_. For clarity, the numerical result for *v* = 0 was displayed only in the case of *N* = 40. : *k* = 0.1, 𝒟 = 0.05 and *L* = 40.

### Chemophoresis engine can recapitulate equi-positioning, directed movement, and pole-to-pole oscillation

First, we considered the motion in a one-dimensional (1D) space (*d* = 1), that is, on a nucleoid matrix along the long cell axis where a plasmid *i*(1 ≤*i* ≤ *M*) was positioned at *x* = *ξ*_*i*_ ∈ [0, *L*] (Fig. 1 B); The Neumann boundary condition was adopted for the RD equation: ∇*u*(0) = ∇*u* (*L*) = 0. To confine the plasmids to the host cell *x* ∈[0, *L*], we placed the reflection walls at *x* = 0 and *x* =*L*. This could be explicitly represented as 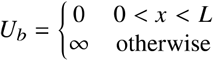. We set *l*_*b*_ *=*0.2 ,*V =*2 *l*_*b*_ *=* 0.4, *m =* 1,*N =*40 through the simulation, and then we examined how the plasmid dynamics change with *χ* := *kN/V*, where *χ* is the normalized maximum rate for ParA-ATP hydrolysis by ParB on each PC. For *M* = 1, the dynamics of *u*(*x*) and the plasmid, as well as the distribution of the plasmid position, are displayed for *χ* = 0.5, 2.5 and 10 in Fig. 1 C. For *χ* = 0.5, the plasmid slightly tends to be localized at the cell center, but it is still dominated by thermal fluctuations (Fig. 1 C1). The plasmid was stably localized at the cell center for *χ* = 2.5 (Fig. 1 C2), whereas for *χ* = 10, it showed directed movement and then reflected at the end walls, resulting in pole-to-pole oscillation (Fig. 1 C3). In general, the plasmid showed directed motion for a larger *χ* (= maximum rate of ParA-ATP hydrolysis). This result was plausible because the larger j generates the sharper gradient of ParA-ATP, leading to the larger chemophoresis force to enable the persistent directed motion of the plasmid.

Similarly, for *M* > 1 cases, plasmid dynamics qualitatively changed among stochastic switching, steady equi-positioning, and directed movement followed by an oscillatory mode as *χ* increased (Fig. S1, and Fig. S2). We then examined how plasmid dynamics switched from a static to an oscillatory mode with increasing *χ*. The switch in plasmid dynamics occurred through a symmetry-breaking transition of the ParA-ATP spatiotemporal pattern. Interestingly, the oscillatory behavior of plasmids did not disrupt time-averaged equi-positioning. Steady multi-modal distribution of plasmids was sustained (Fig. S1, and Fig. S2). As reported previously (2, 3), the regular positioning of plasmids is due to the effective inter-plasmid repulsive interaction derived from the chemophoresis force. The plasmid acting as a sink for ParA-ATP contributed to the formation of a concentration gradient, which increased with the distance from the plasmid. Other plasmids were subjected to the chemophoresis force caused by the gradient in the direction of increasing ParA-ATP concentration so that they were forced away from the former. The former plasmid was also subjected to the chemophoresis force caused by the gradient derived from the latter, resulting in mutual repulsion among the plasmids. The mutual repulsive interaction contributed to the robustness of the positional information generated by the chemophoresis engine. This plasmid separation scenario by such repulsive interactions is consistent with a previous observation (17).

### Chemophoresis engine mathematically validates plasmid surfing on the traveling ParA-ATP wave

In a recent report, (29), Vecchiarelli et al. demonstrated the directed movement of micro-sized beads that mimic plasmids. A theoretical explanation using more realistic model with finit-sized plasmids is needed, as the previous models (3, 37) assumed vanishing-size plasmids; Hence, we introduced the ;_1_-sized plasmids in Eq. 3 and 4, to discuss a possible symmetry-breaking transition. Here, we analytically examined plasmid surfing on the ParA-ATP traveling wave, focusing only on *M* = 1, without thermal fluctuation, for a 1D case under a periodic boundary condition.

Fig. S3 shows simulation results of Eq. 3 and 4 for *χ* = 2.5 (Fig. S3 A) and *χ* = 10 (Fig. S3 B) with *K*_*d*_ = 0.001. In the former case, the plasmid maintained its location, whereas in the latter it traveled on the *u* (*x*) wave and moved unidirectionally. Fig. 2 A shows a steady velocity (=ν) profile of plasmid movement for 0 < *χ* (= :*kN*/(2*l*_*b*_))< 10 and 0 < *ϵ*(= *𝒟N*/(2*l*_*b*_))< 10. In the case without thermal fluctuation, as *χ* and *ϵ* increased, the velocity monotonously increased above a critical curve on the *χ ϵ* plane (Fig. 2 A). Above the curve, the plasmid showed directed motion.

To analytically examine the change in the steady solutions of plasmid dynamics against *χ* values, we simplified Eq. 3 and 4 assuming *u*(*x*) ≫*K*_*d*_ over *x* ∈ [0, *L*]resulting in 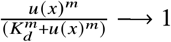 (Supporting Materials and Methods). Furthermore, by introducing a co-moving frame with a space-time coordinate *z* := *x* − *vt*, where *v* is the steady velocity of the plasmid, defining *u*(*x,t*):= *U*(*x* −*vt,t*), and solving the steady-state equation Eq. S3 and S4, we obtained a relationship between *v* and *χ* (Fig. 2 B, green solid line) and the steady-state solution *U*^*st*^ (*z*) (Fig. 2 C, green solid line) (Supporting Materials and Methods). Solutions for directed movement (|*v* |> 0) emerged at *χ* = *χ* _*c*_ 3.1 as a result of a (supercritical) pitchfork bifurcation, whereas the localized solution without motion (*v* = 0) existed over 0 ≤*χ* ≤ 10. Hence, there were three solutions for *χ* > *χ* _*c*_ (Supporting Materials and Methods and Fig. S4 therein). Fig. 2 C shows steady solutions *U*^*st*^ (*z*) for localization (purple) at *χ* = 2.5 and directed movement (green) at *χ* = 10. Note that the plasmid location is fixed at the origin (*z* = 0) on the space-time coordinates (Supporting Materials and Methods). For the case of *v* = 0, the shape of *U*^*st*^ (*z*) was symmetrical and had its minimum at the origin, reaching an equilibrium state of the plasmid location (Fig. 2 C, purple). In contrast, the symmetry of *U*^*st*^ (*z*) was broken for the case of| *v* |> 0, supporting a non-equilibrium traveling wave (Fig. 2 C, green). Interestingly, the minimum of the latter traveling wave was positioned at a location shifted from the origin of the plasmid (Fig. 2 C, inset figure, green), suggesting that the plasmid was “surfing” on the traveling wave. Next, we numerically calculated the steady velocity using Eq. 3 and 4 with *K*_*d*_ = 0.001 over 0 ≤ *χ* ≤ 10 and confirmed the emergence at *χ* = *χ* _*c*_ and the stability over *χ* > *χ* _*c*_ of the solution of the equation Eq. S1 and S2 for the plasmid surfing on the traveling wave (Fig. 2 B, red dots). Therefore, these results demonstrated the existence of solutions for plasmid surfing on the traveling wave of ParA-ATP for a 1D case.

Furthermore, we performed a linear stability analysis of the “surfing-on-wave” pattern using Eq. S3 and S4 against external disturbances, in order to examine if tiny perturbation around the stationary solution was amplified or not as *U*(*z,t*) = *U*^*st*^ (*z*) *e*^*λt*^ δ*U* (*z*), and *z*_ξ_ (*t*) = *e*^*λt*^ *δz*_ξ_, by computing eigenvalues *λ* as a function of *χ* (Fig. S5, Supporting Materials and Methods). Then, the stability of the plasmid-surfing pattern and the instability of the plasmid-localized pattern was confirmed for *χ* > *χ* _*c*_ from the absence of any positive real parts of *λ*, Re [*λ*(*v*)]> 0, *v* ≠ 0 (Fig. S5, Supporting Materials and Methods). If there were any positive parts of eigenvalues for E ≠ 0, the perturbation would be amplified, and then the steady traveling wave solution (Fig. 2 C, green) would be collapsed; however, this was not the case. The results showed that plasmid surfing on the traveling wave emerged through a symmetry-breaking transition at a critical maximum rate of ParA-ATP hydrolysis (=*χ* _*c*_) as a pitchfork bifurcation in dynamic systems theory (Fig. 2 B and Fig. S5).

We also examined *l*_*b*_ dependency of the directed motion of the plasmid for the cases *N* = 10, 20, 30, 40, 50, using the analytical solution Eq. (S7 - S9) (Fig. 2 D). There were no solutions for the directed movement of *N* = 10, whereas, for each of the different values in *N* = 20, 30, 40, 50, the velocity of the directed motion (*v* ≠ 0) was monotonously decreased with *l*_*b*_, and an inverse pitchfork bifurcation occurred at a critical value of *l*_*b*_, resulting in the only solution with *v* = 0 (Fig. 2 D). Next, we numerically calculated *l*_*b*_ dependency of the velocity and confirmed the disappearance of the traveling wave solution (Fig. 2 D, black dot plots). The numerical results tended to deviate from the analytical ones in the ranges of small and large *l*_*b*_, suggesting the breakdown of the approximation *u* (*x*) ≫*K*_*d*_. In general, the directed motion with a larger *l*_*b*_ requires a larger *N*. This is because a plasmid (or bead) with large *l*_*b*_ cannot generate the concentration gradient and the resultant chemohoresis force is not sufficient to realize its self-driven directed motion.

Finally, the above surfing also worked for a two-dimensional (2D) case (Fig. S6 and S7). The simulation results for the 2D case in Eq. 3 and 4 are shown for *χ* = 10 (Fig. S6 A) and *χ* = 50 (Fig. S6 B). In the former case, the cargo maintained its location, and *u*(*r*) had a symmetrical shape (Fig. S7 A), whereas directed motion by surfing on an asymmetrical traveling wave of *u*(*r*) was observed for the latter (Fig. S7 B), just like the 1D case.

## DISCUSSION

In this study, to consider a generalized model of the plasmid partition parABS system, a chemophoresis engine was introduced as a coupled dynamical system among the equations of motion for plasmids and the RD equation for ParA-ATP (Fig. 1 A). In the model, plasmid dynamics switched from static to dynamic mode with an increase in the maximum rate of ATP hydrolysis *χ*. The engine demonstrated equi-positioning, directed movement, and pole-to-pole oscillation, as observed in bacterial cells and *in vitro* experiments (Fig. 1 C, Fig. S1 and S2). Note that despite the plasmids’ oscillatory behavior, the regular positioning distributions were sustained (Fig. S1 and S2) due to an effective inter-plasmid repulsive interaction derived from the chemophoresis force, indicating the robustness of positional information generated by the chemophoresis engine. By simplifying Eq. 3 and 4, and introducing a space-time coordinate, we mathematically showed the existence (Fig. 2, B and C) and the stability (Fig. S5) of the plasmid-surfing pattern. The solution emerged through the symmetry-breaking transition of the ParA-ATP spatiotemporal pattern at a critical *χ*. With an increase of the plasmid size *l*_*b*_, the solution for the directed movement disappeared as a result of an inverse pitchfork bifurcation (Fig. 2 D). We also demonstrated plasmid surfing in a 2D case Fig. S6 and S7).

Although we analyzed the existence and stability of the surfing-on-wave pattern only in a noiseless situation (Fig. 2), plasmids (or cargos) are always subjected to thermal fluctuations in cellular environments. Then, the plasmid motion was described by Langevin equation Eq. 4. Further, we needed to elucidate that the plasmid-surfing-on-traveling-wave pattern remains robust against thermal fluctuations. For the chemophoresis force to act effectively, the force must be larger than the thermal noise, as discussed in a previous report (3). Any force weaker than thermal noise cannot sustain even regular positioning (3). We also examined how equi-positioning of plasmids can overcome thermal noise disturbances in a 1D case (Fig. S1, and S2). We confirmed that plasmid location dynamics shows a transition from stochastic switching to (freezing) steady equi-positioning (Fig. S1, and S2) as j is increased, finally leading to persistent directed motion of the plasmids. This result suggested that the chemophoresis force dominates and directed movement of plasmids can overcome against thermal fluctuation for large hydrolysis rate.

## CONCLUSION

We propose a chemophoresis engine, a general mechano-chemical apparatus driving the self-motion of the intracellular cargo, as a means to elaborate the physical principles of ATPase-driven cargo transport (45–50). The engine is based on 1) a chemophoresis force that allows motion along an increasing ATPase(-ATP) concentration and 2) an enhanced catalytic ATPase hydrolysis at the cargo positions. ATPase-ATP molecules are used as fuel to supply free energy by applying the chemophoresis force along the concentration gradient, whereas cargos generate a concentration gradient by catalyzing the hydrolysis reaction on their surface. Note that each cargo, as a catalyst, does not consume ATP, but only modulates the concentration pattern. Through the coupling and synergy between 1) and 2), directed movement of the cargo is self-organized, showing a “surfing-on-traveling-wave” pattern (Fig. 2 C). The chemophoresis engine is based only on these two general mechanisms and is expected to explain how the transportation of diverse cargos in bacterial and eukaryotic cells is organized. Although we have focused on the gradient generated by the regulation of ATPase, the regulation of the concentration gradient via phosphorylation-dephosphorylation reactions is ubiquitous. Therefore, the chemophoresis engine resulting from the regulation of the hydrolysis of other factors, such as GTPase, should work for a variety of intracellular processes (70–73).

Furthermore, recent studies demonstrated substrate-driven chemotactic behaviors of metabolic enzymes (single-molecule chemotaxis) (74, 75), and discussed a mathematical model to recapitulate experimental results in the subsequent study (76). Interestingly, the authors proposed the exact same thermodynamic mechanism as the chemophoresis force described previously (2, 3). Therefore, we expect that the chemophoresis engine can also be applied to self-chemotactic behaviors even at a single-molecule level even though in the present study, self-chemotaxis is applied to a cargo size ranging from 50 nm to 1’m. However, to describe nanoscopic chemotaxis, we need to extend thermodynamics of chemophoresis to a stochastic one which is valid even under large thermal/chemical fluctuations.

The merits of the chemophoresis engine are as follows: Self-generated chemical gradient for the chemophoresis force to apply; not requiring a large space to maintain the external chemical gradient. The chemophoresis engine can be effective in a moderate space. Therefore, the chemophoresis engine would work for eukaryotic intra-nuclear processes by restricting the mobility of chemicals on a nuclear membrane or a nuclear matrix functioning as a scaffold matrix. We introduced a chemophoresis engine to represent the general mechanism of cargo motion, applying it to plasmid dynamics in a parABS system, leading to its directed motion, equi-positioning, and pole-to-pole oscillation observed in bacterial cells and *in vitro* experiments. Based on the generality of the mechanism and suggestive previous reports (77–79), we can apply the mechanism to other hydrolysis events, RNAs, receptors, and others. We propose the chemophoresis engine as a universal mechanism for general hydrolysis-driven cargo transports in cells.

## APPENDIX A

In this appendix, we discuss the chemical thermodynamics and the corresponding statistical mechanics required for the derivation of the chemophoresis force discussed in Appendix B.

### Rate equation at chemical equilibrium

Consider a macroscopic object, such as organelles or macromolecules that have several binding sites on their surface for chemical reactions to occur. We considered the chemical equilibrium of the reactions at a given temperature *T* and a chemical pool containing a chemical X with a chemical potential *µ m* molecules of X in an environmental solution are attached to binding site B on the bead and form complex Y, as represented by the following reaction: 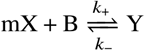. The binding of ligands with a receptor on the membrane is a typical example. The molecular number of the complexes (unbound sites) on the surface is denoted by *N*_*Y*_(*N*_*B*_).. Note that we defined the bead as a macroscopic entity relative to an X molecule. Under a constant site number 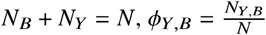 was defined as the volume fraction of Y and B on the bead, respectively. The chemical potential *µ*of the X molecules in the environmental solution was assumed to be constant. The rate equation of the reaction was written as follows: 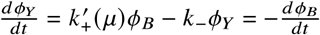 where 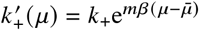 The reaction was assumed to reach chemical equilibrium,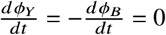. Therefore at equilibrium,

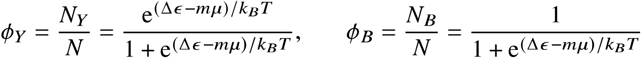

Here 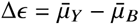. This result was simultaneously satisfied by *N*_*B*_ + *N*_*Y*_= *N* and the condition of the chemical equilibrium *mµ* = *µ*_*Y*_ *µ*_*B*._

### Free energy changes among chemical equilibrium states attached to the chemical pool

Thermodynamics describes transitions among equilibrium states. Under isothermal and fixed surface area conditions, the change in the free energy of the bead can be represented as follows: *dF* (*N*_*Y*_, *N*_*B*_) = *µ*_*Y*_*dN*_*Y*_+*µ*_*B*_*dN*_*B*_ whereas that of the environment is 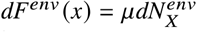 Note that 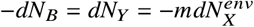 from a stoichiometric relation, *v*_*X*_ = −*m, v*_*B*_ = −1, *v*_*Y*_ = 1 defined by 0 ⇄ *v*_X_X + *v*_B_B + *v*_Y_Y related to the above reaction. Hence the change of the free energy was obtained as follows: *dF* (*N*_*Y*_) = (*µ*_*Y*_−*µ*_*B*_)*dN*_*Y*._. Rewriting *dF* (*N*_*Y*_) using an extent of reaction *θ*, d*θ* = −*dθ*_*B*_ = *dθ*_*Y*._ = −*mdθ*_*X*_ and an affinity *A* of the bead, *A* := ∑ _*i=y, B*_ *v*_*i*_ *μ*_*i*_ = − (*μ*_*Y*_ − *μ*_*B*_) = − *mμ* . From *dθ* = *dθY* = *dN*_*Y*_, *dF*(*θ*) = − *Adθ* (= *mμ dθ*. The affinity of the environment was *A*^*env*^ = − ∑ _*i=y, B*_ *v*_*i*_ *μ*_*i*_ = *mμ*. The value of the total system was *A*^*tot*^ = +*A* ^*env*^ = 0.

Next we considered chemical equilibrium in a chemical pool in the environment. Using the Legendre transform for *θ* using *A*= −(*µ*_*Y*_ − *µ*_*B*_) = −*m*µ*, d* Ω (*A*) := *dF* (*θ*) + *d* (*A *θ*) = *θ* dA* = −*m,d µ*. Here, *θ* = *N*_*Y*_ + *const*. from *d θ* = *d θ*_*Y*._ = *dN*_*Y*._ ,^1^. Therefore,

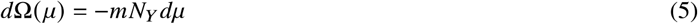

### Statistical mechanics at chemical equilibrium

Next, we considered the corresponding statistical model for the two-state system with Y (bound) and B(unbound) states for the reaction *m* X + B ⇄ Y. The thermodynamic variables were temperature *T*, the affinity of the reaction ,*A* and the extent of reaction *θ*, whereas microscopic variables were the binding energy *ϵ* and binding state *n*. The chemical equilibrium was sustained by a chemical bath containing a chemical X with a chemical potential *µ*, where the equilibrium condition is *A*^*tot*^ = *A*+*A* ^*env*^ = −(*µ*_*Y*_ − *µ*_*B*_) + *mµ* = 0.

If the system was in the bound (unbound) state, the change in its energy was *ϵ* = *ϵ*_*Y*_(*ϵ*_*B*_). For a system with only one binding site, the partition function was obtained by summing the microscopic states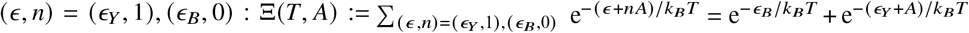. Assuming that the chemical adsorption reactions at the *N* binding sites were independent of each other, the grand potential free energy for the system with # binding sites was written as follows:

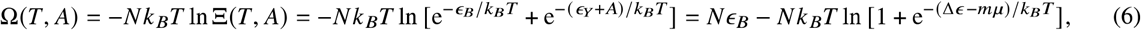

where Δϵ = *ϵ* _*Y*_ − *ϵ* _*B*_. Here we used *A=* − (*µ*_*Y*_ *µ*_*B*_) = *mµ* Therefore, the macroscopic molecular numbers of the bound states *N*_*Y*_ was given by 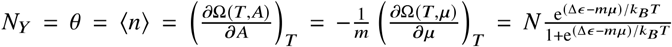 Then, the number of unbound State was 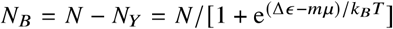. The mixing entropy was also obtained from the free energy: 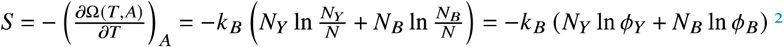.^2^

## APPENDIX B

### Chemophoresis force

We refer to the biological elements defined in Appendix A as beads with several reaction sites to which X molecules attach. The bead was placed at *r* = ***ξ*** and moved in a 3-dimensional space ***r*** *∈* ***R***^*d*^ (*d* = 1, 2, 3). We considered an isothermal process that was homogeneous in a given space at a given temperature *T*, and a chemical pool containing the chemical X with a spatially dependent concentration *x* (***r***) or, equivalently, the corresponding chemical potential *µ* (*r*). This gradient was assumed to be sustained externally. The X molecule was attached to binding site B on the bead and formed a complex Y as represented by the reaction mX **(ξ)**+ ⇄ Y.

The molecular number of the complexes on the bead was denoted by *N*_*Y*_. Note that we defined the bead as a macroscopic entity relative to an X molecule. To consider local equilibrium conditions, the bead was assumed to move sufficiently slowly so that the above reaction was at local chemical equilibrium at the position ***r*** = ***ξ***. In other words, the time scale of the chemical reaction (*τ*_*chem*_) was much smaller than that of the motion of the bead, *τ*_*bead*_ ≫ *τ*_*chem*_ With the assumption that a local equilibrium existed, we applied thermodynamics with spatially dependent thermodynamic variables. The change in the grand potential free energy of the bead was presented by dΩ **(*ξ*)** = *mN*_*Y*_*dµ* (*ξ*) from Eq. 5.

Then, we considered a virtual displacement of the bead. Under an infinitesimal displacement ***ξ → ξ + d ξ***, the change in the grand potential was *d*Ω(***ξ***) = −*mN*_*Y*_*dµ*(*ξ*) = *mN*_*Y*_∇*µ*. (*ξ*) *d ξ*. In other words, the position of the bead, ***ξ***, was adopted as an effective independent variable instead of the chemical potential *µ* (*ξ*). Then, ***ξ*** was used as the work coordinate, and − *mN*_*Y*_***∇****µ*.(***ξ***) was the force exerted on the bead by the external environment, balanced by the force generated by the reservoir of chemical potential distribution *µ* (***r***), which also acts on the bead. Therefore, the reservoir-generated chemical gradient force was represented as follows:

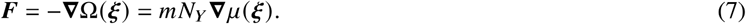

This expression was obtained as described below. Consider a quasi-static infinitesimal displacement *d* ***ξ***. Then, from the change in the grand potential, the maximum work done by the system on the external world through the reservoir is *d* ^*′*^*W =N*_*Y*_***∇****µ*.(***ξ***) *d* ***ξ***. Considering that this work was done by the force exerted by the reservoir on the bead, that is, *d* ^*′*^*W =* ***F***.*d* ***ξ***,, the force formula in Eq.(7). In other words, without an externally applied force, ***ξ*** evolved spontaneously such that Ω (*ξ*) monotonically decreased, that is, *d* Ω(***ξ***) < 0. When we considered an overdamped system, in which the kinetic energy of the bead was negligible, the phenomenological equation of motion was given by 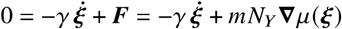, and it was assumed that the friction constant resulting in dissipation was proportional to the velocity with the proportionality constant *γ*.

From the chemical equilibrium condition, 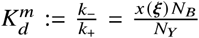, where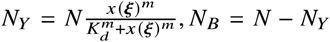. We used the chemical potential of a dilute solution, 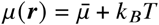, where 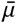 is the standard chemical potential. See Appendix A for details. Finally, the equation of motion was written as follows: 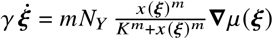 This equation was a general expression for the motion of an element that has a number of binding sites for chemical adsorption under a gradient of chemical potential. We called this motion *chemophoresis* —similar to the nomenclature of typical ‘phoresis’ phenomena. Because of the chemophoresis force, the direction of motion of the bead increased the chemical potential.

In cellular environments, cargo to be modeled as the beads are always subjected to thermal fluctuations, which we included by adding thermal noise ***η***(*t*) and replacing the equation with the Langevin equation, as

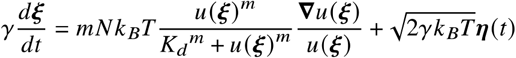

with thermal noise ⟨ **η** (*t*)⟩ = 0 and ⟨ η (*t*) η (*t ′*) ⟩= 2*dγ k*_*B*_*T δ* (*t* − *t′*), and *δ* is the friction constant. For the chemophoresis force to act effectively, the force must be larger than the thermal noise, as discussed in a previous report (3).

## Chemophoresis engine for plasmid motion

We considered a plasmid *i* (1 ≤ *i* ≤ *M*) “ placed into and moving in a *d*-dimensional space ***r*** *∈* ***R***^*d*^ (*d* = 1 or 2) (Fig. 1 A). ParA-ATP was bound to a PC on plasmid 8 at position ***r*** = ***ξ****m* ParA-ATP molecules interacted with ParB, which stimulated ParA ATPase activity at a catalytic rate :; *N* ParB molecules were assumed to be recruited to each PC at ***r*** = ***ξ***. Because ParA could not bind to PC when it was not combined with ATP, free ParA products were released from the PC immediately after ATP hydrolysis. Thus the reaction was written as follows:

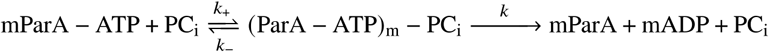

Through this reaction on the PCi at *r* = *ξ*, each plasmid acted as a sink for ParA-ATP and induced a concentration gradient of this protein. Here, we assumed an adiabatic approximation (chemical equilibrium) of the reaction mParA − ATP + PC_i_ ⇄ (ParA - ATP)_*m*_ − PC_i_. Denoting the concentration (chemical potential) of ParA-ATP as *u*(***r***), the time evolution for the ParA-ATP concentration was generally written as follows: *∂tu*(***r***) = -**∇ *J*** (***r***) + ℛ(***r***). Here ***J*** (***r***) is the diffusion flux defined by the Fick law for the chemical potential, 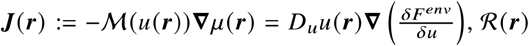 is the reaction term, and *F*^*env*^ [{*u*(***r***)}] is the total free energy of the environment. Considering the free energy of a dilute solution, 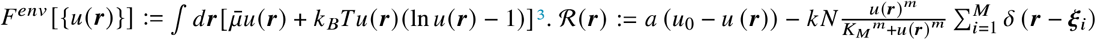 Therefore, the RD equation for the ParA-ATP concentration was given as follows: 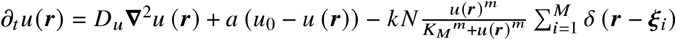 where the first and second terms represent the diffusion of ParA-ATP and its chemical exchange at a constant rate a with the cytoplasmic reservoir (denoted by its concentration *u*_0_), respectively. The last term denotes the inhibition by ParB on *M* PCs. 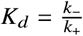 is the dissociation constant, *m* is the Hill coefficient, and δ(***r***) is a delta function representing the hydrolysis reaction point. Without the last term (if *k* = 0), *u*(***r***) reaches a homogenous equilibrium state, *u*(***r***) = *u*_0_. In contrast, the equations of motion for plasmids were given as 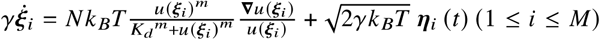 with thermal noise ⟨ **η**_*i*_ (*t*)⟩ = 0 and ⟨ **η**_*i*_ (*t*)· **η**_*j*_ (*t′*)⟩= 2*d γk*_*B*_ *Tδij δ(t −t ′)*,where γ is the friction constant. In these two equations, the size of the plasmid was assumed to be zero (3). To better describe the spatiotemporal profiles of ParA-ATP concentration and directed movement of plasmids, we considered plasmids as a sphere with a radius of *l*_*b*_, and rewrote these equations as 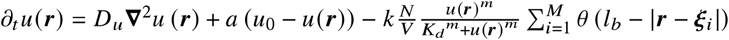, where *V* is the spherical volume of each PC, and *V*=2*l*_*b*_ for *d*= 1 and 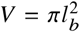 for *d* = 2.*θ*(*r*) is a step function representing the space each PC occupies to describe the hydrolysis reaction space. Only within |*r ξ*_*i*_ |< *l*_*b*_, the reaction occurred. When *l*_*b*_; → 0, *θ* (***r***) / *V* → γ (***r***) ,this equation was reduced to the previous model. By using dimensionless variables, 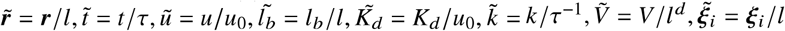 and redefining 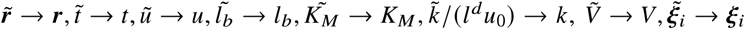 the normalized equation was written as follows:

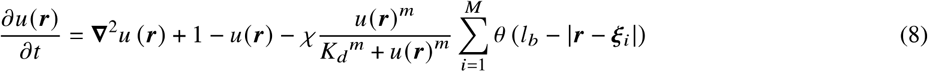

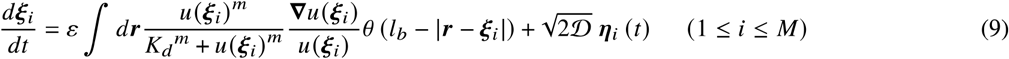

Here, *τ* = *a*^-1^,*l*;^2^ = D_*u*_ *τ*, D_*ξ*_*=kBT*/γ 𝒟 =D_*ξ*_/D_*u*_ is the relative diffusion coefficient of the plasmid to that of ParA-ATP, and ε = 𝒟*N*/*V. χ* = *kN*/*V* is the maximum rate of ParA-ATP hydrolysis by ParB on each PC.

## AUTHOR CONTRIBUTIONS

T.S. performed mathematical modeling and simulation, wrote the manuscript, and was responsible for the coordination of the study; K.K. wrote the manuscript and was responsible for the research project.

## ACKNOWLEDGMENTS

We thank H. Niki, K. Maeshima, A. Kimura, H. Nishimori, A. Awazu, S. Sawai, N. J. Suematsu, S. Nakata, S. Ito, and Y. Okada for their comments. T.S. was supported by JSPS KAKENHI (Grant No. 19H05796).

## SUPPORTING INFORMATION

### SI TEXT

#### Simplified Equations of Chemophoresis Engine for Analytical Examination

In order to examine analytically the change of steady solutions of the plasmid dynamics against the values of *χ* = *kN* / (2*l*_b_) and ε = 𝒟*N* / (2*l*_b_), we simplify Eq. 3 and Eq. 4 in the main text with *M* = 1, without thermal fluctuation, and for a one-dimensional case under a periodic boundary condition. On the assumption of *u*(*x*) ≫ *k*_*d*_ over *x* ∈ [0, *L*] resulting in 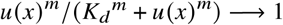, Eq. 3 and Eq.4 are given as:

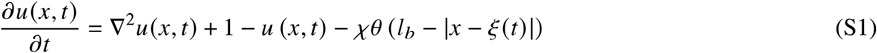

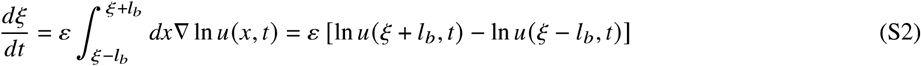

Eq. S2 gives a well-defined representation for the non-equilibrium condition under chemophoresis force on the plasmid through the (normalized) chemical potential difference from end to end Δ _*μ*_:= ln *u* (*ξ* + *l*_*b*_) *−*ln *u* (*ξ − l*_*b*_), which is self-generated by the plasmid.

By introducing a spacetime coordinate *z= x −vt*, where *v* is a constant velocity of the frame, and defining *u* (*x, t*) ≔ *U* (*x − vt, t*), Eq. S1 and S2 are rewritten as

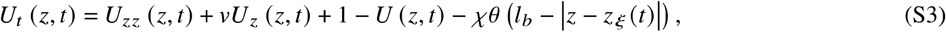

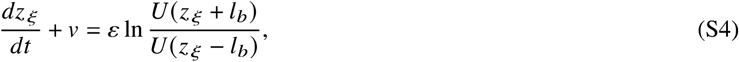

where *U*_*z*_ shows a partial differential for *z* and *z*_*ξ*_ (*t*) = *ξ* (*t*) − *vt* is a spacetime position of the plasmid. By considering a steady state of the plasmid’s position in *z* space, the coordinate represents a co-moving frame, where *v* corresponds to a steady velocity of the plasmid. A steady state equation of Eq. S3 and S4 is given from *U*_*t*_ = 0 and ż *ξ* = 0 as:

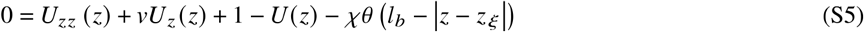

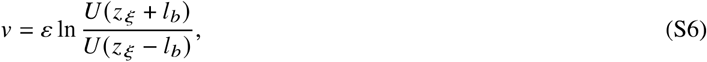

Here, at the steady state, *z ξ* = 0 can be set without losing generality. By solving Eq. S5 and S6, the steady state solution *U*^*st*^ (*z*) and *v* is obtained as:

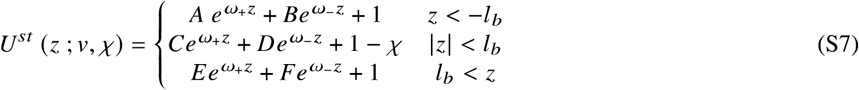

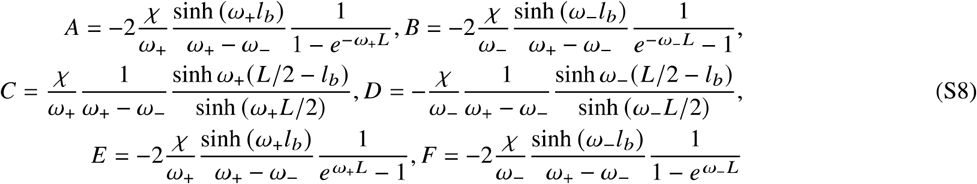

and 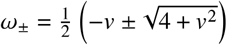. Finally by calculating *v* from the following equation numerically, we obtain a relationship between *v* and *χ* (Fig. 2 B, green solid line in the main text):

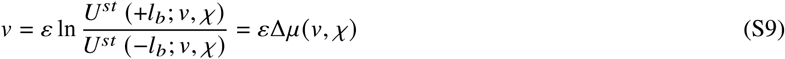

Eq. S9 gives a self-consistent equation for *v*; The chemical potential difference Δ *μ* (*v, χ*) determined by the bead’s velocity *v* generates the chemophoresis force acting on itself, resulting in its steady velocity *v*. Eq. S9 for E shows a pitch-fork bifurcation at *χ* = *χ* _*c*_ ∼ 3.1, and has three solutions at *χ* > *χ* _*c*_ (Fig. S4).

#### Linear Stability Analysis of “Plasmid Surfing on Traveling Wave” Solution

We performed linear stability analysis of surfing-on-wave solution for Eq. S3 and S4 against external disturbances, in order to examine if tiny perturbation around the stationary solution is amplified or not, as *U*(*z, t*) = *U*^*st*^ (*z*) + *e* ^*λt*^ *δU*(*z*), and 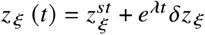, by computing eigenvalues *λ* as a function of *χ* (Fig. S5). Substituting the perturbation into Eq. S3 and S4, we obtain by using the steady solution Eq. S5 and S6 with 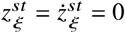:

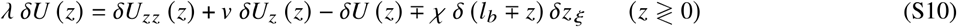

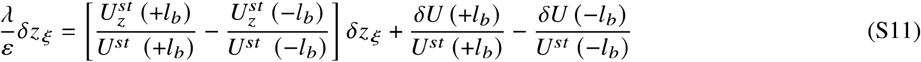

Furthermore, by setting a jump condition (1), the following formulas are given as:

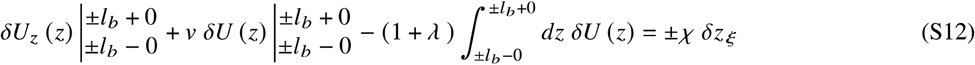

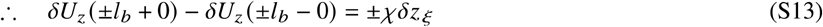

By solving Eq. S10 with the jump condition Eq. S13, the two boundary conditions *δU*(−*L*/2) = *δU* (*L*/2), *δUz*(−*L*/2) = *δUz* (*L*/2), and the two continuity conditions of *δU* (z) at *z* = ±*l*_*b*_, *δU* (±*l*_*b*_ − 0) = *δU* (±*l*_*b*_ + *0)*, the solution *δU* (z) is obtained as a function of *δz*_*ξ*_ :

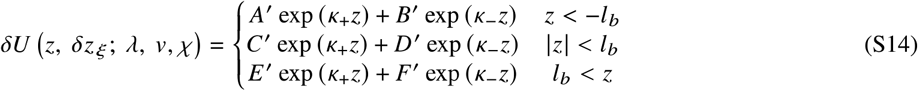

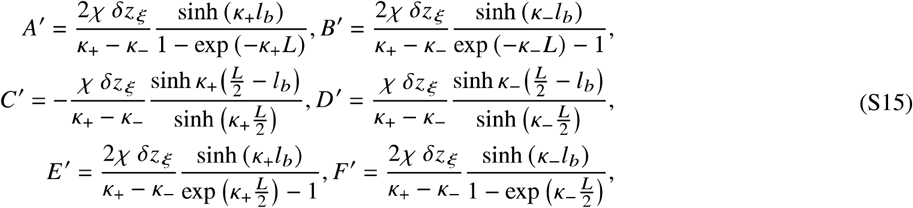

and 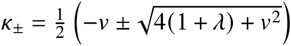. From Eq. S11 and S14, we get an eigenvalue equation for *λ*:

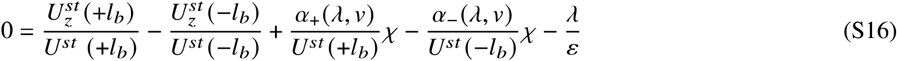

Here, 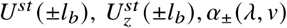 and are given as:

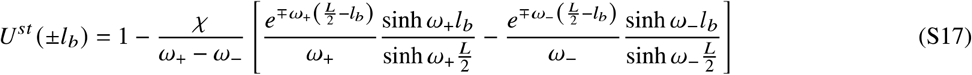

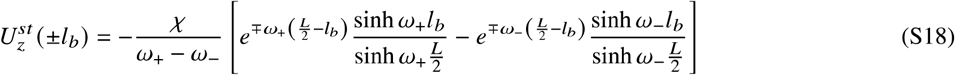

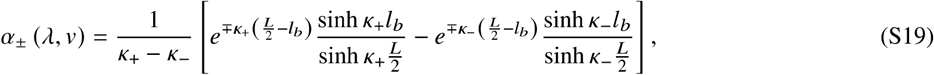

where *λ* satisfying with the eigenvalue equation S16 are complex numbers in general. Then, the stability of the plasmid-surfing solution (Fig 2C, green) and the instability of the plasmid-localized solution are confirmed for *χ* > 1 from the absence of max Re [*λ* (*v*)] > 0 for *v* ≠ 0 and the existence of max Re [_(0)] > 0, respectively (Fig. S5). If Eq. S16 with E < 0 contained any positive real parameter Re [*λ* (*v*)] > 0, the perturbation would be amplified and then the steady traveling wave solution (Fig. 2C, green) would be collapsed. However, this is not the case; max Re [*λ* (*v*)] = 0 is obtained for *χ* larger than the bifurcation point *χ* _*c*_ ∼ 3.1 for *v* ≠ 0 (Fig. S5). Here, Re [*λ* (*v*)]= 0 for the surfing solution (Fig 2C, green) corresponds to Goldstone mode in physics, where the perturbation is neither amplified nor decayed, but only shifts the solution along the I axis due to the translational invariance (See (2) for details of the Goldstone mode studied mathematically in reaction-diffusion systems).

On the other hand, the localized solution is destabilized at *χ* = *χ*_*c*_ and Re [*λ* (0)] turns out to be positive, whereas it regains stability at *χ* larger than *χ* ∼ 5.9 (Fig. S5). However, such localized solution cannot be numerically realized for *χ* > 5.5 because *U*^st^ (0) becomes negative and the solution is unphysical at *χ* ∼ 5.5 (Fig. S5, inset) as a result of breaking the approximation u(x) ≫ *k*_*d*_; Therefore, a stable localized solution does not exist for *χ* ≳ 5.5.

Finally, these results show that plasmid surfing on the traveling wave emerges through a symmetry-breaking transition at a critical reaction rate of ParA-ATP hydrolysis (=*χ*) as a pitch-fork bifurcation in dynamical systems theory (Fig. 2B and S5).

## SUPPORTING FIGURES

**Figure S1:**
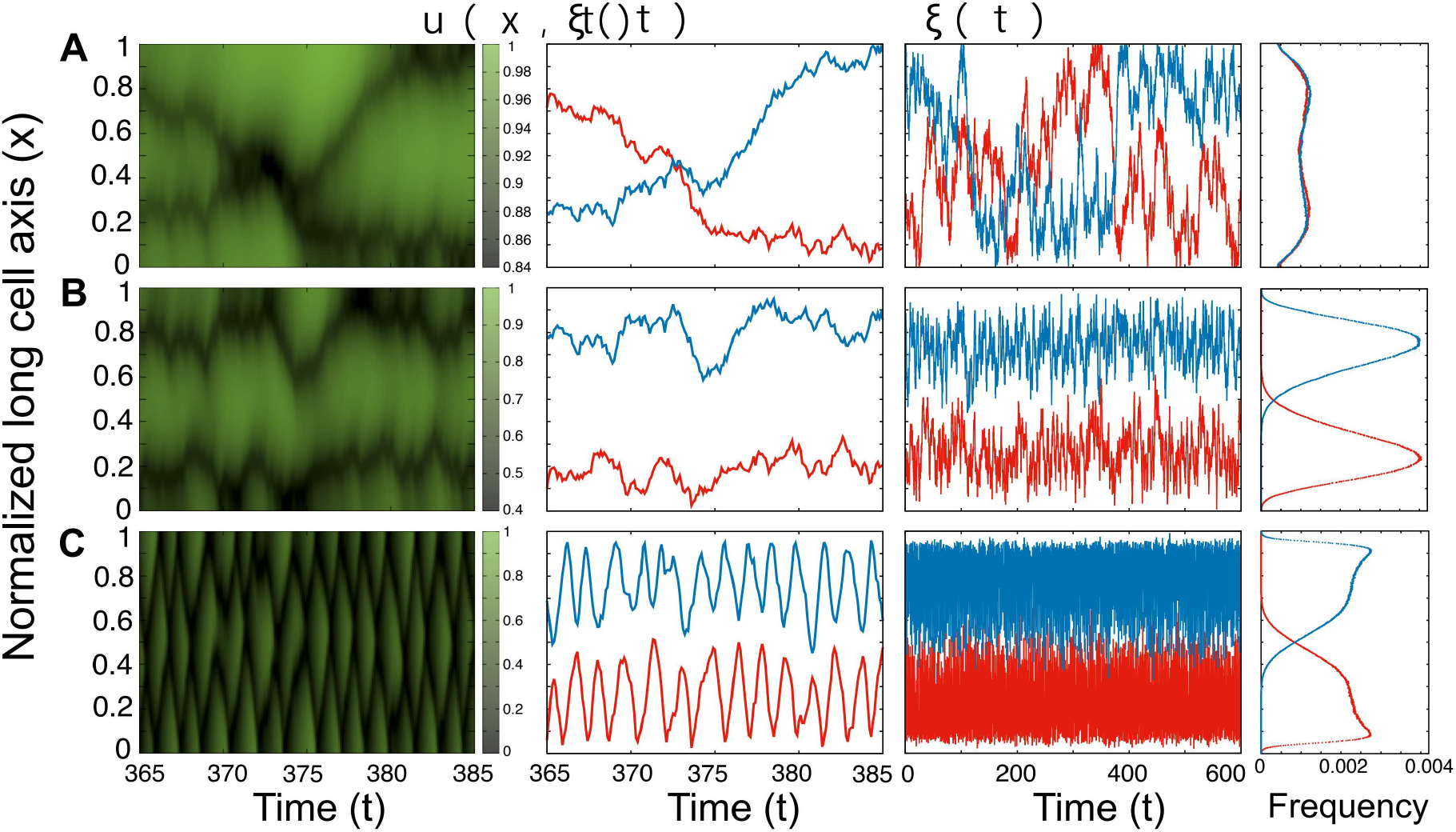
Plasmid location dynamics for M = 2. The dynamics change among stochastic switching, steady equi-positioning, and directed movement followed by oscillatory mode as *χ* increases among *χ* = 0.5 (A), *χ* = 2.5 (B), and *χ* = 10 (C) (two inner figures). The corresponding ParA-ATP pattern dynamics also change among stochastic switching, steady equi-positioning, and oscillatory waves (left). The oscillatory behavior of plasmids does not disrupt time-averaged equi-positioning. Steady multi-modal distributions of plasmids are sustained (Compare B right and C, right). k_*d*_ = 0.1, ε = 5, and *L* = 5. The distributions (right) were generated using 10^7^ samples over 10^5^ time step.

**Figure S2:**
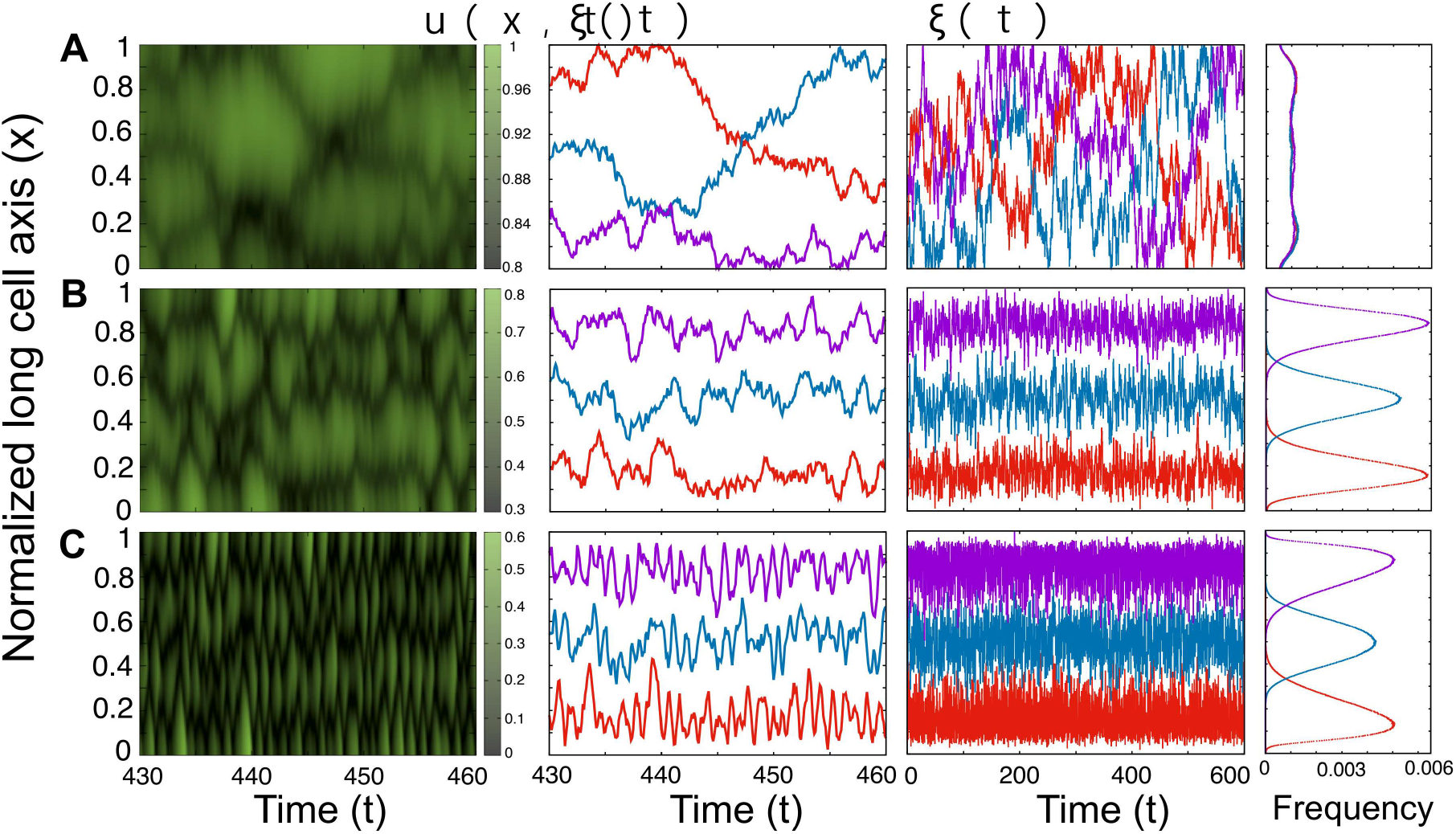
Plasmid location dynamics for *M* = 3. The dynamics change among stochastic switching, steady equi-positioning, and directed movement followed by oscillatory mode as *χ* increases among *χ* = 0.5 (A), *χ* = 2.5 (B), and *χ* = 10(C) (two inner figures). The corresponding ParA-ATP pattern dynamics also change among stochastic switching, steady equi-positioning, and oscillatory waves (left). The oscillatory behavior of plasmids does not disrupt time-averaged equi-positioning. Steady multi-modal distributions of plasmids are sustained (Compare B right and C, right). *k*_*d*_ = 0.1, *ε* = 5, and *L* = 5. The distributions (right) were generated using 10^7^ samples over 10^5^ time step.

**Figure S3:**
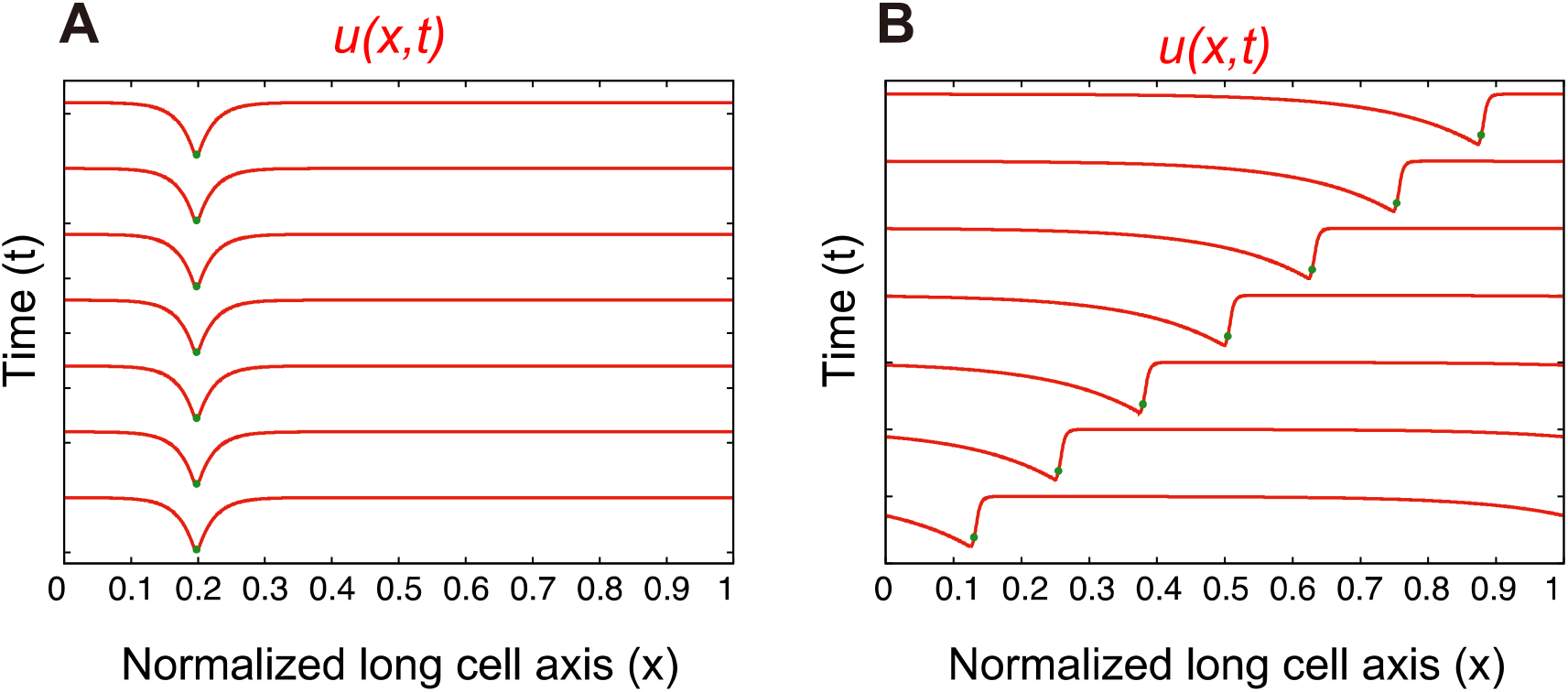
Simulation results of Eq. 3 and 4 (see SI Text) for *χ* = 2.5 (A) and *χ* = 10 (B) with *k*_*d*_= 0.001. The red lines show the spatial pattern of *u*(*x, t*) (=ParA-ATP) for each C. Green dots show a plasmid location for each C. In the former case (A), the plasmid maintains its location, whereas it surfs on the traveling wave of D G, C and moves unidirectionally in the latter (B). These results were compared with the analytical results in the main text and in Fig. 2. *ε* = 5 and *L* = 40.

**Figure S4:**
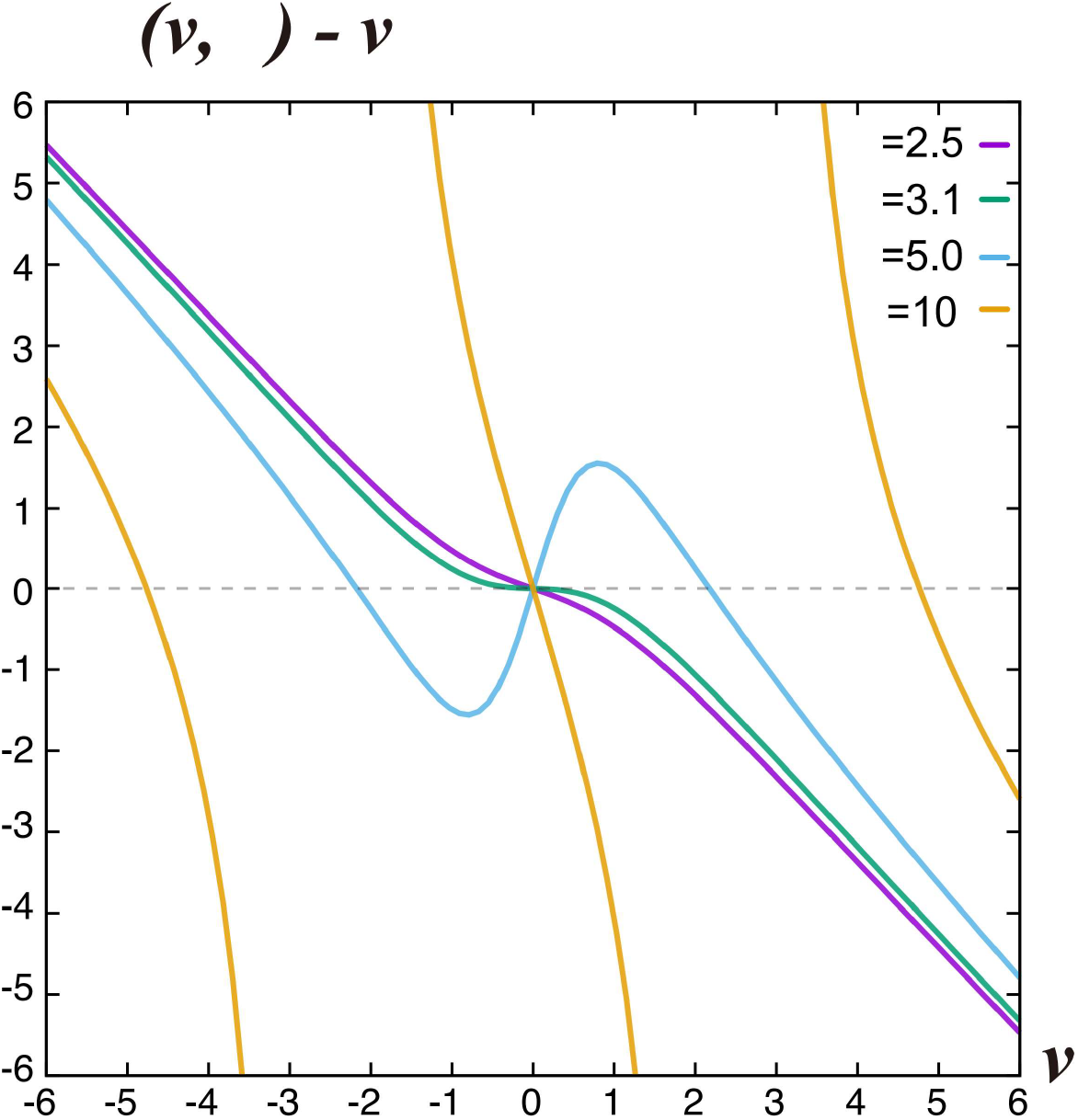
Analytical solution of *ε Δ μ* (*v, χ*) − *v* for a visualization of the self-consistent equation Eq. S9. Eq. S9 shows a pitch-fork bifurcation at *χ* = χ_*c*_∼ 3.1, and has three solutions for *χ* > *χ*_*c*_. *ε* = 5 and *L* = 40.

**Figure S5:**
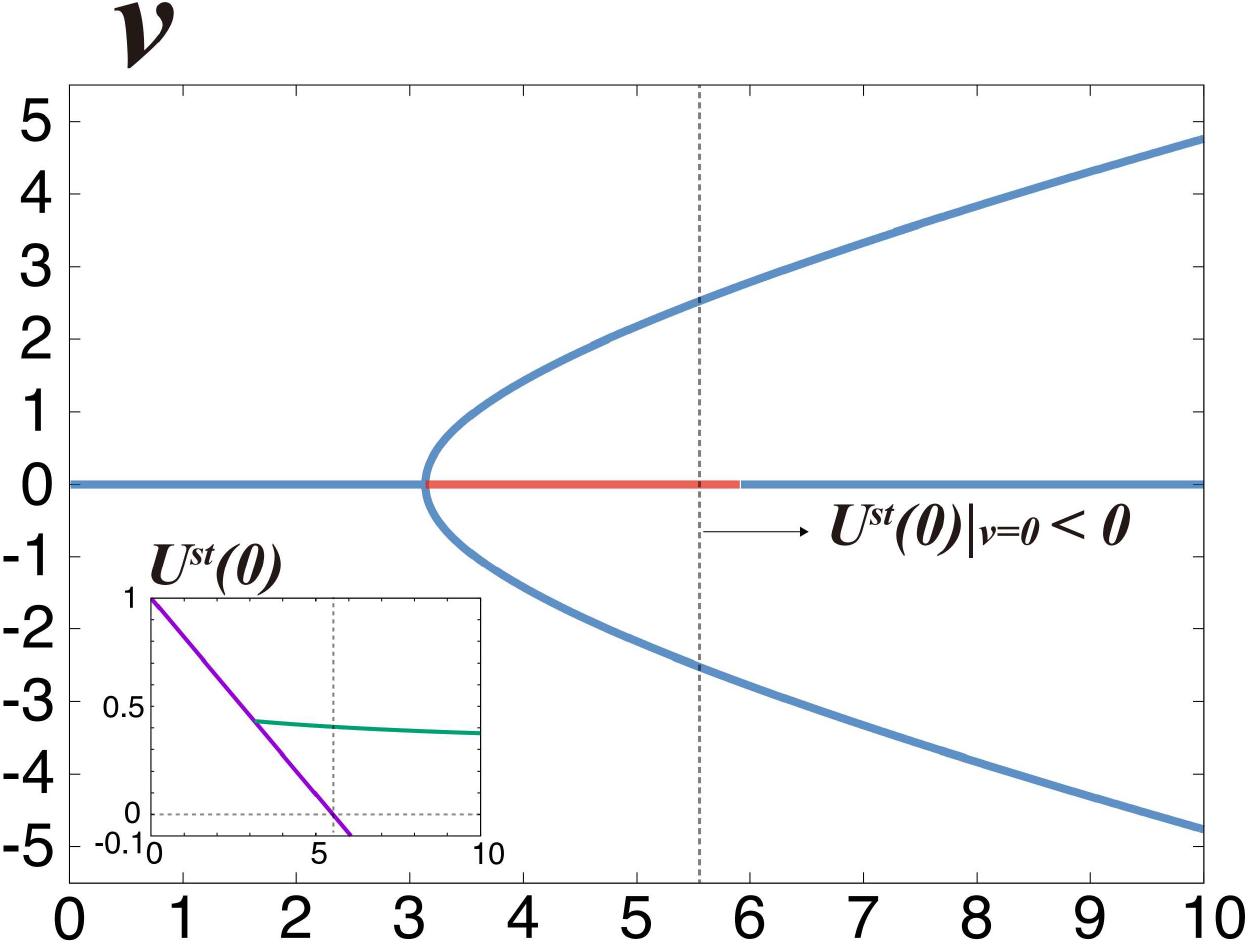
Linear stability analysis of the surfing-on-wave pattern *U*^*st*^ (*z*) and 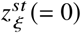 against external disturbances. From tiny perturbations around the stationary solution *U* (*z, t*) = *Ust* (*z*) + *e*λ*t δU* (*z*), and 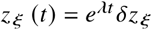, the eigenvalues *λ* were computed as a function of χ. The stability of the plasmid-surfing pattern (blue line) and the instability of the plasmid-localized solution (red line) was confirmed for *χ* > χ_*c*_ in the absence of any positive real parameters of *λ*, Re [*λ*(*v*)] > 0, *v* ≠ 0. It seems to regain stability at *χ* larger than *χ* 5.9. However, such localized solution cannot be numerically realized for *χ* > 5.5 because *U*^*st*^(0) becomes negative and the solution is unphysical at *χ* ∼ 5.5 (inset) as a result of breaking the approximation *u*(*x*) ≫ *k*_*d*_ Therefore, the stable localized solution does not exist for *χ* ≳ 5.5. *N* = 40, and *L* = 40.

**Figure S6:**
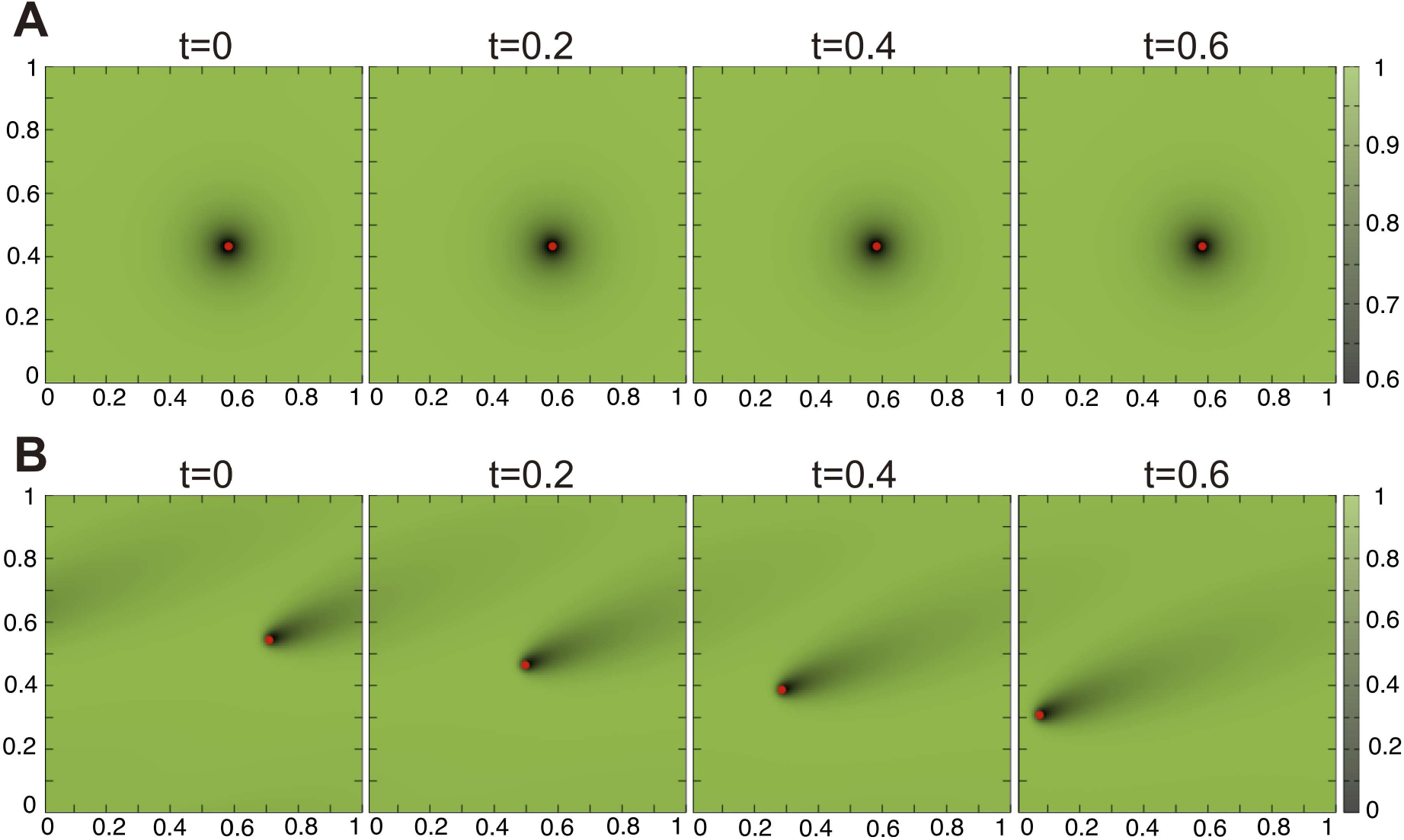
Simulation results for a two-dimensional (2D) case of Eq. 3 and 4 (see SI Text). Successive snapshots are shown for *χ* = 10 (A) and *χ* = 50 (B). In the former case (A), the plasmid (red circle) maintains its location at a minimum symmetrical shape of u(r, t) (=ParA-ATP, green scale), whereas it moves unidirectionally with an asymmetrical pattern of D(A, C) in the latter (B). *k*_*d*_ = 0.001, ;*l*_b_ = 0.2, ε = 5, and *L*^2^ = 10 × 10.

**Figure S7:**
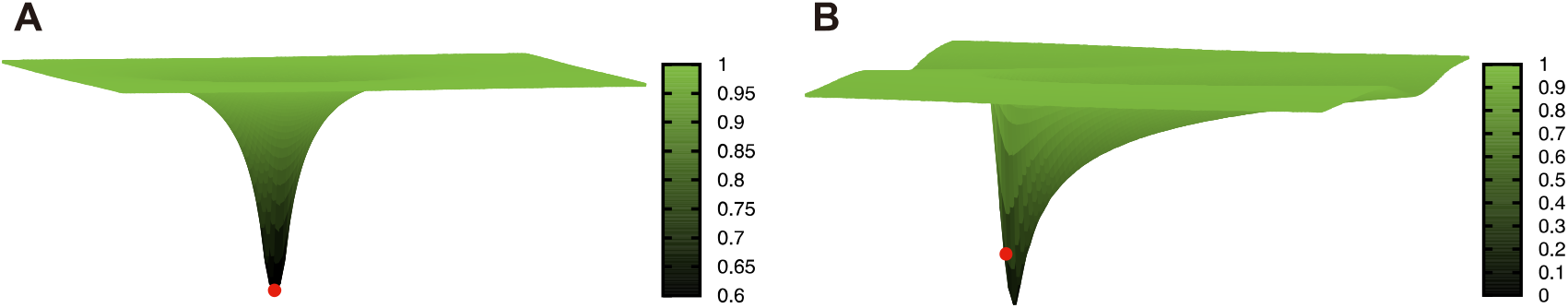
Simulation results for a two-dimensional (2D) case of Eq. 2 and 3 (see SI Text). Snapshots are shown for *χ* = 10 (A) and *χ* = 50 (B). In the former case (A), *u*(*r, t*) (=ParA-ATP, green scale) is symmetrical, and the plasmid (red circle) is located at its minimum. In contrast, for the latter (B), the symmetry of *u*(*r, t*) is broken, indicative of a traveling wave. The minimum of the asymmetrical D(A, C) is positioned at a location shifted from where the plasmid lies, suggesting that the plasmid in “surfing” on the traveling wave *k*_*d*_ = 0.001, *l*_b_ = 0.2, ε = 5, and *L*^2^ = 10 × 10.

## Footnotes

1 *const* = 0 from the initial condition (*θ* _Y._ ,N_Y._) = (0, 0).

2 We used 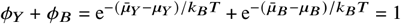 in the calculation.

3 If we studied the effect of phase separation, we could introduce *F*^*env*^ leading to the Cahn-Hilliard equation.

## REFERENCES

1. Turing, A., 1952. Philosophical the royal biological transqfctions society sciences. Phil. Trans. R. Soc. Lond. B 237:37–72.

2. Sugawara, T., 2010. Chemophoresis in a cell. PhD thesis, The University of Tokyo, Tokyo, Japan.

3. Sugawara, T., and K. Kaneko, 2011. Chemophoresis as a driving force for intracellular organization: Theory and application to plasmid partitioning. Biophysics 7:77–88.

4. Yang, H. W., S. R. Collins, and T. Meyer, 2016. Locally excitable Cdc42 signals steer cells during chemotaxis. Nature cell biology 18:191–201.

5. Das, M., T. Drake, D. J. Wiley, P. Buchwald, D. Vavylonis, and F. Verde, 2012. Oscillatory dynamics of Cdc42 GTPase in the control of polarized growth. Science 337:239–243.

6. Yuan, X.-b., M. Jin, X. Xu, Y.-q. Song, C.-p. Wu, M.-m. Poo, and S. Duan, 2003. Signalling and crosstalk of Rho GTPases in mediating axon guidance. Nature cell biology 5:38–45.

7. Niki, H., and S. Hiraga, 1997. Subcellular distribution of actively partitioning F plasmid during the cell division cycle in E. coli. Cell 90:951–957.

8. Gordon, G. S., D. Sitnikov, C. D. Webb, A. Teleman, A. Straight, R. Losick, A. W. Murray, and A. Wright, 1997. Chromosome and low copy plasmid segregation in E. coli: visual evidence for distinct mechanisms. Cell 90:1113–1121.

9. Mohl, D. A., and J. W. Gober, 1997. Cell cycle–dependent polar localization of chromosome partitioning proteins in Caulobacter crescentus. Cell 88:675–684.

10. Li, Y., and S. Austin, 2002. The P1 plasmid is segregated to daughter cells by a ‘capture and ejection’mechanism coordinated with Escherichia coli cell division. Molecular microbiology 46:63–74.

11. Li, Y., A. Dabrazhynetskaya, B. Youngren, and S. Austin, 2004. The role of Par proteins in the active segregation of the P1 plasmid. Molecular microbiology 53:93–102.

12. Gordon, S., J. Rech, D. Lane, and A. Wright, 2004. Kinetics of plasmid segregation in Escherichia coli. Molecular microbiology 51:461–469.

13. Toro, E., S.-H. Hong, H. H. McAdams, and L. Shapiro, 2008. Caulobacter requires a dedicated mechanism to initiate chromosome segregation. Proceedings of the National Academy of Sciences 105:15435–15440.

14. Fogel, M. A., and M. K. Waldor, 2006. A dynamic, mitotic-like mechanism for bacterial chromosome segregation. Genes & development 20:3269–3282.

15. Schofield, W. B., H. C. Lim, and C. Jacobs-Wagner, 2010. Cell cycle coordination and regulation of bacterial chromosome segregation dynamics by polarly localized proteins. The EMBO journal 29:3068–3081.

16. Shebelut, C. W., J. M. Guberman, S. Van Teeffelen, A. A. Yakhnina, and Z. Gitai, 2010. Caulobacter chromosome segregation is an ordered multistep process. Proceedings of the National Academy of Sciences 107:14194–14198.

17. Sengupta, M., H. J. Nielsen, B. Youngren, and S. Austin, 2010. P1 plasmid segregation: accurate redistribution by dynamic plasmid pairing and separation. Journal of bacteriology 192:1175–1183.

18. Lenz, P., and L. Søgaard-Andersen, 2011. Temporal and spatial oscillations in bacteria. Nature Reviews Microbiology 9:565–577.

19. Howard, M., and K. Gerdes, 2010. What is the mechanism of ParA-mediated DNA movement? Molecular microbiology 78:9–12.

20. Vecchiarelli, A. G., K. Mizuuchi, and B. E. Funnell, 2012. Surfing biological surfaces: exploiting the nucleoid for partition and transport in bacteria. Molecular microbiology 86:513–523.

21. Vecchiarelli, A. G., L. C. Hwang, and K. Mizuuchi, 2013. Cell-free study of F plasmid partition provides evidence for cargo transport by a diffusion-ratchet mechanism. Proceedings of the National Academy of Sciences 110:E1390–E1397.

22. Hwang, L. C., A. G. Vecchiarelli, Y.-W. Han, M. Mizuuchi, Y. Harada, B. E. Funnell, and K. Mizuuchi, 2013. ParA-mediated plasmid partition driven by protein pattern self-organization. The EMBO journal 32:1238–1249.

23. Kiekebusch, D., and M. Thanbichler, 2014. Spatiotemporal organization of microbial cells by protein concentration gradients. Trends in Microbiology 22:65–73.

24. Hatano, T., Y. Yamaichi, and H. Niki, 2007. Oscillating focus of SopA associated with filamentous structure guides partitioning of F plasmid. Molecular microbiology 64:1198–1213.

25. Hatano, T., and H. Niki, 2010. Partitioning of P1 plasmids by gradual distribution of the ATPase ParA. Molecular microbiology 78:1182–1198.

26. Treuner-Lange, A., and L. Søgaard-Andersen, 2014. Regulation of cell polarity in bacteria. Journal of Cell Biology 206:7–17.

27. Vecchiarelli, A. G., K. C. Neuman, and K. Mizuuchi, 2014. A propagating ATPase gradient drives transport of surface-confined cellular cargo. Proceedings of the National Academy of Sciences 111:4880–4885.

28. Kiekebusch, D., and M. Thanbichler, 2014. Plasmid segregation by a moving ATPase gradient. Proceedings of the National Academy of Sciences 111:4741–4742.

29. Vecchiarelli, A. G., Y. Seol, K. C. Neuman, and K. Mizuuchi, 2014. A moving ParA gradient on the nucleoid directs subcellular cargo transport via a chemophoresis force. BioArchitecture 4:154–159.

30. Lim, H. C., I. V. Surovtsev, B. G. Beltran, F. Huang, J. Bewersdorf, and C. Jacobs-Wagner, 2014. Evidence for a DNA-relay mechanism in ParABS-mediated chromosome segregation. Elife 3:e02758.

31. Vecchiarelli, A. G., J. A. Taylor, and K. Mizuuchi, 2015. Reconstituting ParA/ParB-mediated transport of DNA cargo. Methods in cell biology 128:243–269.

32. Hu, L., A. G. Vecchiarelli, K. Mizuuchi, K. C. Neuman, and J. Liu, 2015. Directed and persistent movement arises from mechanochemistry of the ParA/ParB system. Proceedings of the National Academy of Sciences 112:E7055–E7064.

33. Ietswaart, R., F. Szardenings, K. Gerdes, and M. Howard, 2014. Competing ParA structures space bacterial plasmids equally over the nucleoid. PLoS computational biology 10:e1004009.

34. Surovtsev, I. V., H. C. Lim, and C. Jacobs-Wagner, 2016. The slow mobility of the ParA partitioning protein underlies its steady-state patterning in Caulobacter. Biophysical journal 110:2790–2799.

35. Surovtsev, I. V., M. Campos, and C. Jacobs-Wagner, 2016. DNA-relay mechanism is sufficient to explain ParA-dependent intracellular transport and patterning of single and multiple cargos. Proceedings of the National Academy of Sciences 113:E7268–E7276.

36. Hu, L., A. G. Vecchiarelli, K. Mizuuchi, K. C. Neuman, and J. Liu, 2017. Brownian ratchet mechanism for faithful segregation of low-copy-number plasmids. Biophysical journal 112:1489–1502.

37. Walter, J.-C., J. Dorignac, V. Lorman, J. Rech, J.-Y. Bouet, M. Nollmann, J. Palmeri, A. Parmeggiani, and F. Geniet, 2017. Surfing on protein waves: proteophoresis as a mechanism for bacterial genome partitioning. Physical Review Letters 119:028101.

38. Hanauer, C., S. Bergeler, E. Frey, and C. P. Broedersz, 2021. Theory of Active Intracellular Transport by DNA-relaying. Physical Review Letters 127:138101.

39. Surovtsev, I. V., and C. Jacobs-Wagner, 2018. Subcellular organization: a critical feature of bacterial cell replication. Cell 172:1271–1293.

40. Merino-Salomón, A., L. Babl, and P. Schwille, 2021. Self-organized protein patterns: The MinCDE and ParABS systems. Current Opinion in Cell Biology 72:106–115.

41. Yamaichi, Y., and H. Niki, 2000. Active segregation by the Bacillus subtilis partitioning system in Escherichia coli. Proceedings of the National Academy of Sciences 97:14656–14661.

42. Bignell, C., and C. M. Thomas, 2001. The bacterial ParA-ParB partitioning proteins. Journal of biotechnology 91:1–34.

43. Lutkenhaus, J., 2012. The ParA/MinD family puts things in their place. Trends in microbiology 20:411–418.

44. MacCready, J. S., J. L. Basalla, and A. G. Vecchiarelli, 2020. Origin and evolution of carboxysome positioning systems in cyanobacteria. Molecular biology and evolution 37:1434–1451.

45. Savage, D. F., B. Afonso, A. H. Chen, and P. A. Silver, 2010. Spatially ordered dynamics of the bacterial carbon fixation machinery. Science 327:1258–1261.

46. MacCready, J. S., P. Hakim, E. J. Young, L. Hu, J. Liu, K. W. Osteryoung, A. G. Vecchiarelli, and D. C. Ducat, 2018. Protein gradients on the nucleoid position the carbon-fixing organelles of cyanobacteria. Elife 7:e39723.

47. Borden, J. S., and D. F. Savage, 2021. New discoveries expand possibilities for carboxysome engineering. Current Opinion in Microbiology 61:58–66.

48. Thompson, S. R., G. H. Wadhams, and J. P. Armitage, 2006. The positioning of cytoplasmic protein clusters in bacteria. Proceedings of the National Academy of Sciences 103:8209–8214.

49. Roberts, M. A., G. H. Wadhams, K. A. Hadfield, S. Tickner, and J. P. Armitage, 2012. ParA-like protein uses nonspecific chromosomal DNA binding to partition protein complexes. Proceedings of the National Academy of Sciences 109:6698–6703.

50. Ringgaard, S., M. Zepeda-Rivera, X. Wu, K. Schirner, B. M. Davis, and M. K. Waldor, 2014. ParP prevents dissociation of CheA from chemotactic signaling arrays and tethers them to a polar anchor. Proceedings of the National Academy of Sciences 111:E255–E264.

51. Thanbichler, M., and L. Shapiro, 2006. MipZ, a spatial regulator coordinating chromosome segregation with cell division in Caulobacter. Cell 126:147–162.

52. Kiekebusch, D., K. A. Michie, L.-O. Essen, J. Löwe, and M. Thanbichler, 2012. Localized dimerization and nucleoid binding drive gradient formation by the bacterial cell division inhibitor MipZ. Molecular cell 46:245–259.

53. Raskin, D. M., and P. A. de Boer, 1999. MinDE-dependent pole-to-pole oscillation of division inhibitor MinC in Escherichia coli. Journal of bacteriology 181:6419–6424.

54. Lutkenhaus, J., 2007. Assembly dynamics of the bacterial MinCDE system and spatial regulation of the Z ring. Annu. Rev. Biochem. 76:539–562.

55. Loose, M., K. Kruse, and P. Schwille, 2011. Protein self-organization: lessons from the min system. Annual review of biophysics 40:315–336.

56. Loose, M., E. Fischer-Friedrich, J. Ries, K. Kruse, and P. Schwille, 2008. Spatial regulators for bacterial cell division self-organize into surface waves in vitro. Science 320:789–792.

57. Loose, M., E. Fischer-Friedrich, C. Herold, K. Kruse, and P. Schwille, 2011. Min protein patterns emerge from rapid rebinding and membrane interaction of MinE. Nature structural & molecular biology 18:577.

58. Vecchiarelli, A. G., M. Li, M. Mizuuchi, L. C. Hwang, Y. Seol, K. C. Neuman, and K. Mizuuchi, 2016. Membrane-bound MinDE complex acts as a toggle switch that drives Min oscillation coupled to cytoplasmic depletion of MinD. Proceedings of the National Academy of Sciences 113:E1479–E1488.

59. Vecchiarelli, A. G., M. Li, M. Mizuuchi, V. Ivanov, and K. Mizuuchi, 2017. MinE recruits, stabilizes, releases, and inhibits MinD interactions with membrane to drive oscillation. bioRxiv doi.org/10.1101/109637.

60. Hunding, A., G. Ebersbach, and K. Gerdes, 2003. A mechanism for ParB-dependent waves of ParA, a protein related to DNA segregation during cell division in prokaryotes. Journal of molecular biology 329:35–43.

61. Ringgaard, S., J. van Zon, M. Howard, and K. Gerdes, 2009. Movement and equipositioning of plasmids by ParA filament disassembly. Proceedings of the National Academy of Sciences 106:19369–19374.

62. Ebersbach, G., and K. Gerdes, 2001. The double par locus of virulence factor pB171: DNA segregation is correlated with oscillation of ParA. Proceedings of the National Academy of Sciences 98:15078–15083.

63. Ebersbach, G., and K. Gerdes, 2004. Bacterial mitosis: partitioning protein ParA oscillates in spiral-shaped structures and positions plasmids at mid-cell. Molecular microbiology 52:385–398.

64. Ebersbach, G., S. Ringgaard, J. Møller-Jensen, Q. Wang, D. J. Sherratt, and K. Gerdes, 2006. Regular cellular distribution of plasmids by oscillating and filament-forming ParA ATPase of plasmid pB171. Molecular microbiology 61:1428–1442.

65. Bonny, M., E. Fischer-Friedrich, M. Loose, P. Schwille, and K. Kruse, 2013. Membrane binding of MinE allows for a comprehensive description of Min-protein pattern formation. PLoS Comput Biol 9:e1003347.

66. Kruse, K., M. Howard, and W. Margolin, 2007. An experimentalist’s guide to computational modelling of the Min system. Molecular microbiology 63:1279–1284.

67. Meinhardt, H., and P. A. de Boer, 2001. Pattern formation in Escherichia coli: a model for the pole-to-pole oscillations of Min proteins and the localization of the division site. Proceedings of the National Academy of Sciences 98:14202–14207.

68. Kerr, R. A., H. Levine, T. J. Sejnowski, and W.-J. Rappel, 2006. Division accuracy in a stochastic model of Min oscillations in Escherichia coli. Proceedings of the National Academy of Sciences 103:347–352.

69. Howard, M., and A. D. Rutenberg, 2003. Pattern formation inside bacteria: fluctuations due to the low copy number of proteins. Physical Review Letters 90:128102.

70. Bendezú, F. O., and S. G. Martin, 2013. Cdc42 explores the cell periphery for mate selection in fission yeast. Current biology 23:42–47.

71. Dyer, J. M., N. S. Savage, M. Jin, T. R. Zyla, T. C. Elston, and D. J. Lew, 2013. Tracking shallow chemical gradients by actin-driven wandering of the polarization site. Current biology 23:32–41.

72. Arkowitz, R. A., 2013. Cell polarity: wanderful exploration in yeast sex. Current Biology 23:R10–R12.

73. Ghose, D., K. Jacobs, S. Ramirez, T. Elston, and D. Lew, 2021. Chemotactic movement of a polarity site enables yeast cells to find their mates. Proceedings of the National Academy of Sciences 118:e2025445118.

74. Zhao, X., and A. Sen, 2019. Metabolon formation by chemotaxis. Methods in enzymology 617:45–62.

75. Zhao, X., K. Gentile, F. Mohajerani, and A. Sen, 2018. Powering motion with enzymes. Accounts of chemical research 51:2373–2381.

76. Mohajerani, F., X. Zhao, A. Somasundar, D. Velegol, and A. Sen, 2018. A theory of enzyme chemotaxis: from experiments to modeling. Biochemistry 57:6256–6263.

77. Yehl, K., A. Mugler, S. Vivek, Y. Liu, Y. Zhang, M. Fan, E. R. Weeks, and K. Salaita, 2016. High-speed DNA-based rolling motors powered by RNase H. Nature nanotechnology 11:184–190.

78. Blanchard, A. T., A. S. Bazrafshan, J. Yi, J. T. Eisman, K. M. Yehl, T. Bian, A. Mugler, and K. Salaita, 2019. Highly polyvalent DNA motors generate 100+ pN of force via autochemophoresis. Nano letters 19:6977–6986.

79. Sakai, T., S. I. Nishimura, T. Naito, and M. Saito, 2017. Influenza A virus hemagglutinin and neuraminidase act as novel motile machinery. Scientific reports 7:1–11.

## REFERENCES

1. Nagayama, M., S. Nakata, Y. Doi, and Y. Hayashima, 2004. A theoretical and experimental study on the unidirectional motion of a camphor disk. Physica D: Nonlinear Phenomena 194:151–165.

2. Purwins, H.-G., H. Bödeker, and A. Liehr, 2005. Dissipative solitons in reaction-diffusion systems. In Dissipative solitons, Springer, 267–308.

